# The Maculalactone Biosynthetic Gene Cluster, a Cryptic Furanolide Pathway Revealed in *Nodularia* sp. NIES-3585

**DOI:** 10.1101/2025.02.26.640319

**Authors:** Paul M. D’Agostino, Roberta R. de Castro, Valdet Uka, Kumar Saurav, Yelyzaveta Kriukova, Tobias M. Milzarek, Angela Sester, Maria Paula Schneider, Tobias A. M. Gulder

## Abstract

Cyanobacteria have long been recognised as a prolific source of bioactive natural products (NPs). Among these are the furanolides, a structurally diverse class of compounds first discovered in the 1980’s. Furanolides are characterised by a γ -butyrolactone core bearing aromatic or aliphatic substituents at the α- and β -positions and an aromatic substituent at the γ -position. Recent advances in understanding the genetic basis of furanolide biosynthesis has enabled genome mining approaches to discover related cryptic furanolide biosynthetic gene clusters (BGCs).

In this work, we identified and cloned a cryptic BGC (15.5 kb) from *Nodularia* sp. NIES-3585 using the Direct Pathway Cloning (DiPaC) strategy and heterologously expressed it in *E. coli* BAP1. Through isolation and structural elucidation, we characterised the known compounds maculalactone B and deoxyenhygrolide A and discovered the novel analogue maculalactone N, featuring a 4-hydroxyphenyl substituent at theβposition.

Application of Global Natural Product Social Molecular Networking (GNPS) analysis of high-resolution LCMS data enabled the identification of 25 maculalactone-related molecules. Further, a MS/MS fragmentation rationale for furanolides was developed and used to probe for maculalactone-like molecules that were too low in abundance for isolation. The fragmentation analysis suggests theβ substituent displays remarkable diversity, accommodating aromatic, aliphatic, or indole moieties. Additionally, structural diversity appears through various hydroxylation patterns. These results demonstrate the substrate promiscuity of the maculalactone biosynthetic enzymes and their capacity to generate considerable structural diversity, while highlighting DiPaC as an effective strategy to access cyanobacterial NPs from cryptic BGCs.

## 1. Introduction

Cyanobacteria are recognised as producers of structurally intriguing bioactive natural products (NPs).^1,2^ Among the NPs found in cyanobacteria, furanolides represent a distinct class, characterised by a γ-butyrolactone core structure (**1**) with aromatic or aliphatic substituents at the α- andβpositions, and a conserved aromatic substituent at the γ-position (Figure 1). Cyanobacterial furanolides include the first chlorinated NP discovered from a freshwater cyanobacterium and potent phytotoxin cyanobacterin (**2**),^3,4^ the nostoclides (e.g., **3**),^5,6^ and the maculalactones (e.g., **4-5**).^7–9^

**Figure 1:**
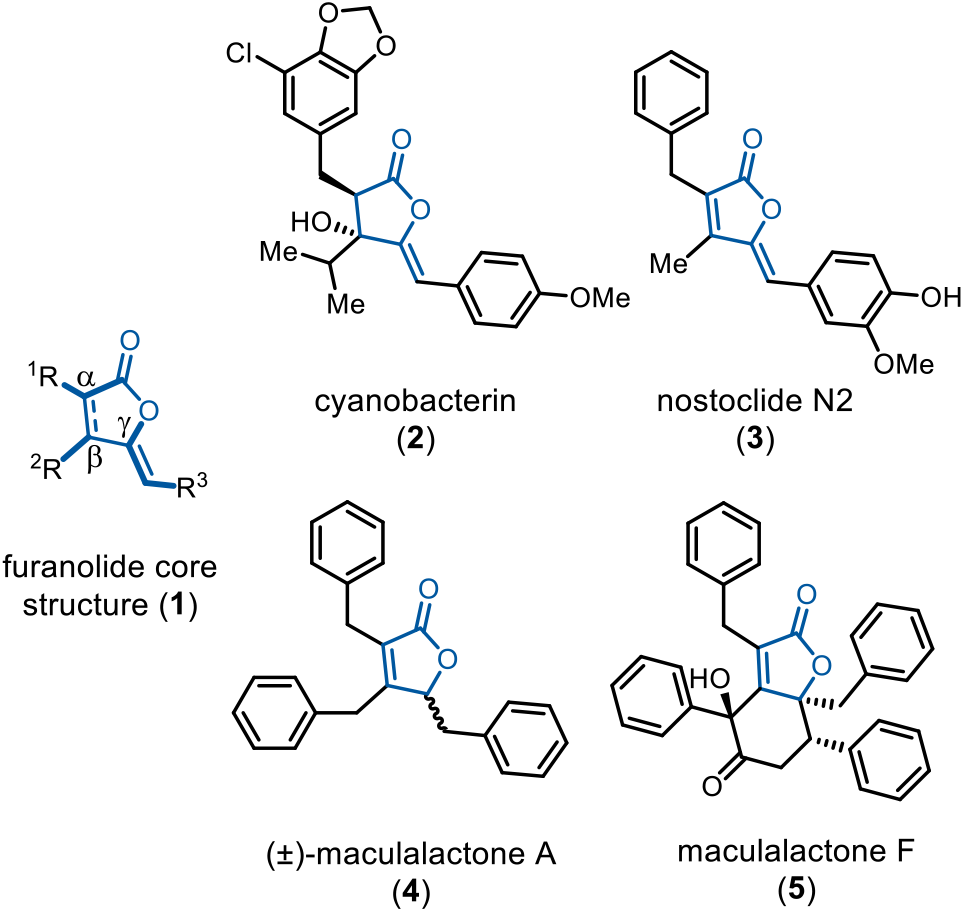
Cyanobacterial furanolides. The γ-butyrolactone core structure **1** (blue) is consistent across the majority of furanolides. The α- andβ positions (R^1^ and R^2^, respectively) consist of aliphatic or aromatic substituents whilst the γ-position (R^3^) retains a conserved aromatic substituent. Representative furanolides include cyanobacterin (**2**), nostoclide N2 (**3**), and the maculalactones, with analogues such as (±)-maculalactone A (**4**) and maculalactone F (**5**).

The maculalactones, exclusively isolated from the marine cyanobacterium *Kyrtuthrix maculans*, collected from rocks at several locations along Hong Kong’s shores, represent the largest and most structurally complex subgroup of furanolides.^8–11^ To date, approximately 14 maculalactone analogues have been reported, exhibiting remarkable structural diversity including tribenzyl-(e.g., **4**) or dibenzyldiphenyl variants (e.g., **5**), all bearing aromatic substituents on the maculalactone core (Figure 1).^8–12^ Furthermore, a racemic maculalactone A ((±)-**4**) was shown to possess antifouling activity against marine herbivores and marine settlers.^7^

Our group recently provided key insights into furanolide biosynthesis by identifying and characterising the cyanobacterin (*cyb*) biosynthetic gene cluster (BGC).^13^ Briefly, biosynthesis occurs through an acyloin condensation reaction catalysed by the thiamine pyrophosphate-dependent (TPP) enzyme CybE, which utilises two α-keto acids to form the first *C*-*C* bond between the α- andβpositions of the furanolide core. The ammonia lyase CybB deaminates L-tyrosine to form 4-coumaric acid, which is subsequently activated to 4-coumaroyl-CoA by the CoA-ligase CybC. The furanolide synthase CybF catalyses a cascade to fuse the acyloin intermediate with 4-coumaroyl-CoA, involving decarboxylative *O*-acylation followed by a Morita–Baylis–Hillman (MBH) reaction and a final 1,4-hydride shift that facilitates late-stage adjustment of the oxidative state, resulting in two additional *C-C* bonds and formation of **1**, common across the furanolide family of NPs.^13^

During our ongoing identification of cryptic furanolide BGCs throughout cyanobacteria, comparative genomics of *Nodularia* sp. NIES-3585 revealed a cryptic BGC encoding all four signature furanolide forming biosynthetic enzymes. Enabled by our Direct Pathway Cloning (DiPaC)^13–22^ strategy for BGC capture and expression, we investigated the cryptic pathway using heterologous expression. Herein, we detail the characterisation of this BGC through integrated molecular, biochemical, and analytical approaches. This work established the maculalactone (*mac*) BGC as the genetic basis for tribenzyl-type maculalactone biosynthesis, the structural elucidation of a novel compound, in addition to the detection of multiple related compounds through molecular networking and MS/MS fragmentation analysis.

## 2. Results and Discussion

### 2.1 Bioinformatic Analysis

The core cyanobacterin biosynthetic protein sequences (CybBCEF) were used to screen the genome of *Nodularia* sp. NIES-3585. This analysis revealed a putative BGC spanning approximately 15.5 kb comprising 11 genes (Figure 2A; Supporting Table S1). This region encoded all four core furanolide biosynthetic genes, consistent with *cyb* and nostoclide N (*ncl*) BGCs, alongside additional genes predicted to be involved in precursor supply.^5,13^ These included 3-deoxy-D-arabinoheptulosonate 7-phosphate (DAHP) synthase (*macA*) and chorismate mutase (*macM*) – both involved in biosynthesis of aromatic amino acids – and a tyrosinase (*macK*), predicted to perform *ortho*-hydroxylation of tyrosine. Further, several open reading frames within the *mac* BGC could not be assigned clear functions, including genes encoding two hypothetical proteins (*orf1* and *orf2*), a NADH-flavin reductase (*orf3*) and a phycocyanobilin:ferredoxin oxidoreductase (*orf4*). The overall composition of the identified candidate BGC strongly suggested it represents a furanolide biosynthetic pathway.

**Figure 2:**
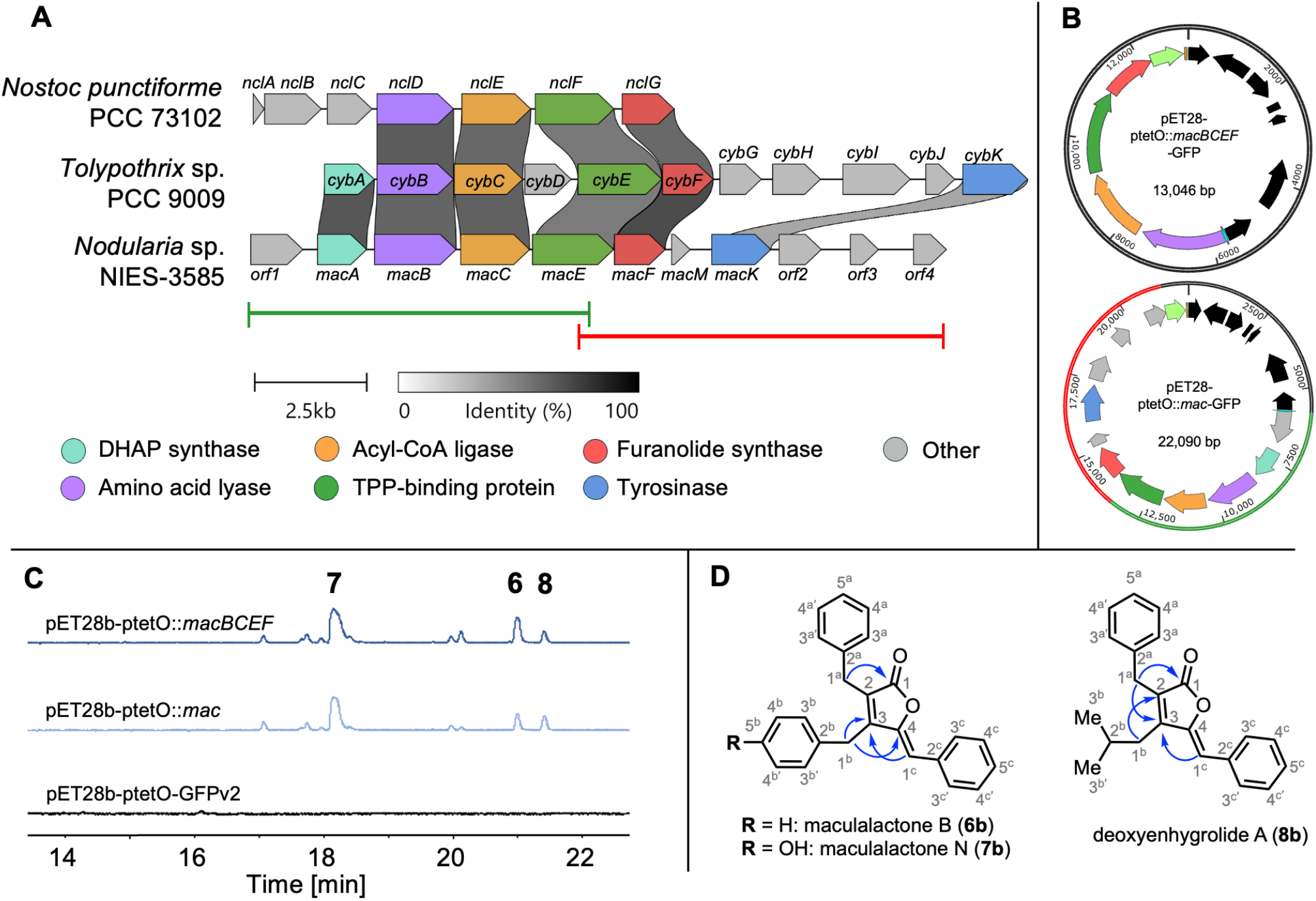
The *mac* BGC and isolated products. **A)** Comparative analysis of the *mac* BGC with the homologous furanolide BGCs, *cyb* and *ncl*.^5,13^ Green and red horizontal bars denote PCR amplicon boundaries used for the DiPaC cloning strategy. **B)** pET28b-ptetO::*macBCEF* and pET28b-ptetO::*mac* plasmid maps. **C)** HPLC-UV chromatograms (330_nm_) comparing organic extracts of cell pellets from experiments with the two expression plasmids to the empty expression vector control. The three compounds selected for isolation and structural characterisation are numbered **6-8. D)** Structures with key HMBC correlations (blue arrows) used for structure elucidation of the isolated compounds **6-8** (for full NMR data, cf. Supporting Table S2 and Supporting Figures S26-S40).

### 2.2 Heterologous Expression, Metabolomic Profiling, and Structure Elucidation

Two separate expression plasmids were constructed to experimentally validate the cryptic BGC and investigate potential roles of additional genes for which no homologues are found in the *cyb* pathway (Figure 2A and 2B). The first construct, pET28b-ptetO::*mac*, harboured the entire BGC in its native arrangement whereas the second construct, pET28b-ptetO::*macBCEF*, contained only the four core furanolide biosynthetic genes (hereafter referred to as *mac* and *macBCEF*, respectively). Following heterologous expression of both plasmids in *E. coli* BAP1, HPLC analysis revealed the production of additional compounds not observed in the control strain harbouring the empty vector backbone (Figure 2C). However, no differences between *mac* and *macBCEF* were observed.

To comprehensively explore the metabolic changes induced by BGC expression, LC-HRMS data from *mac, macBCEF*, and the vector control expression experiments were subjected to multivariate statistical analysis using PCA and PLS-DA.^23,24^ PCA revealed clear separation of both *mac* and *macBCEF* from the control along PC1, whereas PC2 and PC3 showed only minor differences between the two expression constructs (Supporting Figures S1 and S2). PLS-DA and a variable importance projections (VIP) plot corroborated these trends, showing that expressions with *mac* and *macBCEF* were metabolically similar to each other but substantially distinct from that of the vector backbone control (Supporting Figures S3-S5).

Having established the production of new compounds, we proceeded to isolate and elucidate the structures of the three major products (Figure 2C). LC-HRMS analysis of each compound revealed *m/z* values of 353.15 [M+H]^+^, 369.15 [M+H]^+^, and 319.17 [M+H]^+^ with calculated molecular formulae of C_25_H_21_O_2_^+^, C_25_H_21_O_3_^+^, and C_22_H_23_O_2_^+^ for compounds **6, 7**, and **8**, respectively (Table 1). ^1^H and ^13^C NMR analysis confirmed 25 carbons for **6** and **7**, and 22 carbons for **8**, including a characteristic ester signal in each compound (δ_C_ 170.5; 170.6; 170.2) and at least two aromatic phenyl systems consistent with the maculalactone family.^10,12^

**Table 1:**
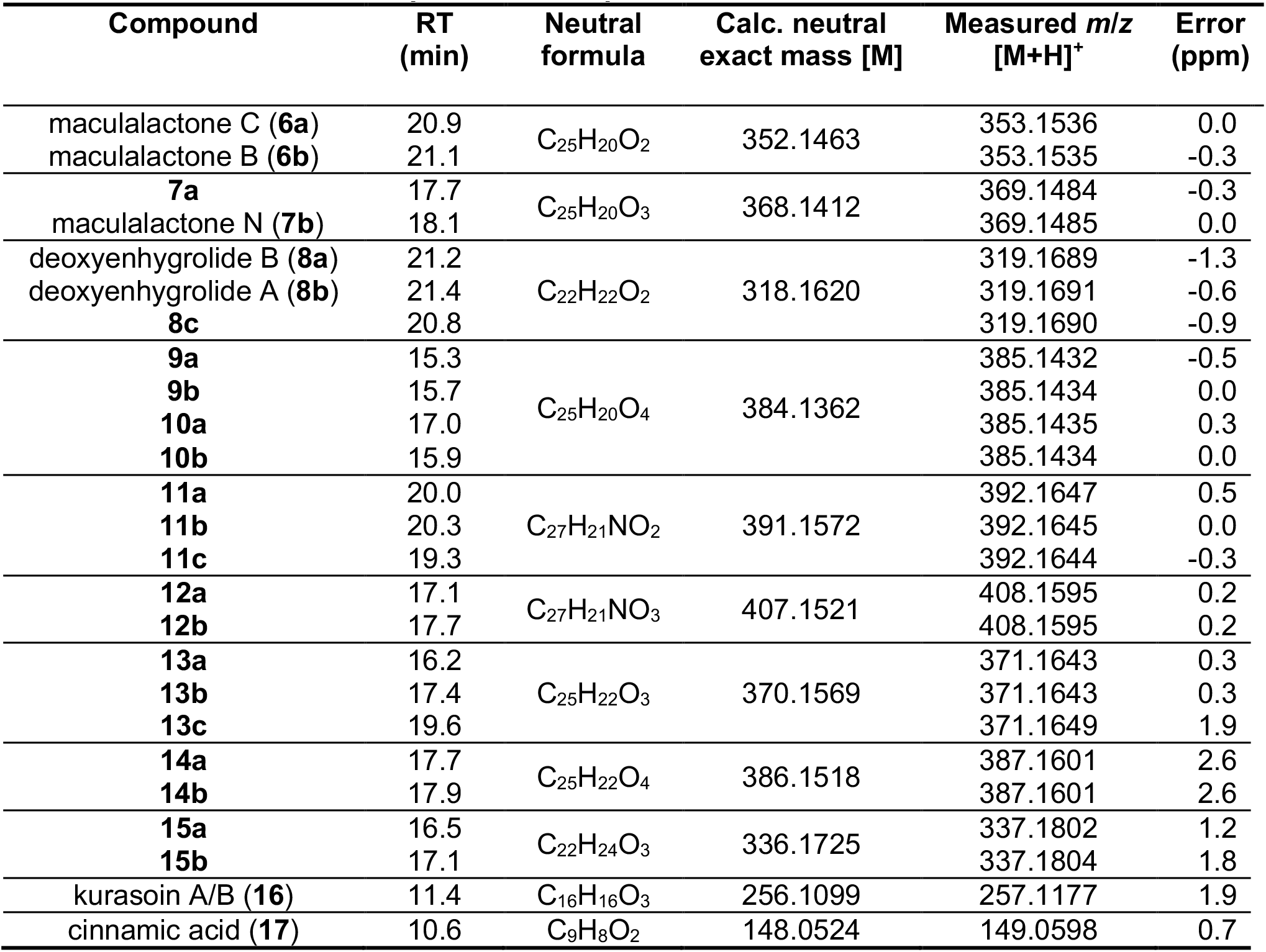
Isolated and MS/MS-predicted compounds.

Compound **6** was confirmed to be maculalactone B/C identified as a mixture of interconverting (*E/Z*)-isomers at positions C4 and C1^c^ (Figure 2D). The conversion of isomers was further probed and due to purity restrictions, we focused on the isolation of compound **6b** and monitored conversion over time by LC-HRMS. Over several days, the more thermodynamically stable *Z*-isomer (**6b**) slowly converted to the less favourable *E*-isomer (**6a**), consistent with synthetically produced **6**^7^ as well as other furanolide NPs including enhygrolide^25^ and precyanobacterin I^13^ (Supporting Figure S6).

The NMR data of compound **7** were highly similar to those of **6**, containing almost identical aromatic substituents. The higher mass (+16 Da) corresponds to an additional oxygen atom and together with characteristic aromatic signals (δ_C_ 154.7; δ_H_ 6.97 (d, 8.5), 6.75 (d, 8.6)) indicates the presence of a 4-hydroxyphenyl moiety (Supporting Table S2). The HMBC correlation between the CH_2_ function (C1^a^ or C1^b^) and the aromatic CH (C4^a^ or C4^b^) next to the hydroxyl group clearly proves that the OH group is not located in the γ-substituted aromatic ring as it would require a similar correlation with the olefinic CH bridge (C1^c^), which was not observed. A strong HMBC correlation of the CH_2_ function at 3.73 ppm to the ester carbonyl (C1, 170.6 ppm) indicated the aromatic substituent devoid of the phenol function to be located at the α-position of **7**. This was further corroborated based on two assignments: (i) the near-exact NMR shifts (Supporting Table S2) when comparing the α-aromatic ring of **6** versus **7**, both indicating a non-hydroxylated α-substituent and (ii) the MS/MS fragmentation pattern of furanolides, in which the α-substituent is always lost first by fragmentation, as observed during in-depth LCMS/MS analyses of a broad range of cyanobacterin- and maculalactone-like furanolides derived of enzymatic synthesis.^26,27^ Thus, a clear assignment of the hydroxyl group on the aromatic ring at the furanolide β -position could be assigned (Supporting Figure S10). Compound **7** was the major product in all expression experiments. This suggests cinnamoyl-CoA (α-position), 4-hydroxyphenylpyruvate (β-position), and phenylpyruvate (γ-position) as the preferred substrates for maculalactone biosynthesis. Our work represents the first report of this NP, which was named maculalactone N (**7**).

Compound **8** differs from **6** and **7** due to the presence of an aliphatic substituent at theβposition. The lower molecular mass (-34 Da relative to **6**), together with the aliphatic NMR signals (δ_H_ 2.32 (d, 7.2), 1.20 (hept, 6.9), 0.46 (d, 6.7)) indicated replacement of an aromatic substituent with an alkyl residue. The coupling pattern, together with the COSY correlations (H1^b^ to H2^b^ to H3^b/b’^), identified the aliphatic residue as an isobutyl group derived from 4-methyl-2-oxopentanoic acid. HMBC correlations (C1^b^ to C2 and C1^b^ to C3) located this substituent at the β-position (Supporting Table S2). Similar isobutyl substituents were observed in *E. coli* extracts and *in vitro* assays of cyanobacterin biosynthetic enzymes, although a clear preference for 3-methyl-2-oxopentanoic acid was reported.^13^ Comparison of **8** to known furanolides revealed the compound to correspond to deoxyenhygrolide A, which was initially reported from the marine myxobacterium *Enhygromyxa* sp. SNB-1.^28^ However, this is the first report of **8** from a cyanobacterium. Compounds **6-8** were submitted for antibiotic activity testing against a range of Gram-positive and Gram-negative bacteria (Supporting Table S3) including more susceptible strains such as *E. coli* Δ*tolC*. However, neither of the three compounds exhibited detectable antibiotic activity.

### 2.3 A Furanolide MS/MS Fragmentation Rationale and Molecular Networking Lead to the Identification of Low Abundance Maculalactone-Like Analogues

To better understand the fragmentation behaviour of the furanolides, the MS/MS spectrum of the already known compound anhydrocyanobacterin within the GNPS library was analysed to generate a putative furanolide fragmentation rationale (detailed description provided in Supporting Text S1; Figure S8). An important characteristic of furanolide fragmentation appears to be the initial cleavage of the α-substituent leading to a strong fragment ion being composed of the furanolide core and both β- and γ-substituents. This step allows for the exact mass of the α-substituent (observed as Da lost) as well as theβ and γ-substituents to be calculated. The initial ion containing theβ and γ-substituent is subsequently further fragmented to often allow assignment of substituents at theβ and γ-positions. Importantly, these patterns were consistently observed in MS/MS spectra of cyanobacterin- and maculalactone-like furanolide derivatives produced both *in vivo* and chemo-enzymatically.^26,27,29^ The fragmentation rationale was applied to assign fragment ions of the three NMR elucidated compounds **6-8** (detailed description Supporting Text S2; Supporting Figure S9-S11).

LCMS/MS data of organic extracts of *in vivo* heterologous expression experiments was used to build a molecular network using the GNPS platform^30^ and a network of nodes that included the parent ions of **6-8** was identified (Figure 3; Supporting Figure S7). A total of 25 structures, including **6-8** and the predicted maculalactone precursors kurasoin A/B (**16**) and cinnamic acid (**17**), were putatively assigned as maculalactone-related NPs based solely on extensive MS/MS fragment ion annotations (Table 1, **9-17**, Supporting Figure S9-S25). The identified maculalactones could be classified into three structural groups based on their substituent patterns and modifications. The first and most abundant group comprised analogues with aromatic substituents at the α-,β and γ-positions, indicating substrate preferences of the maculalactone core biosynthetic enzymes. The compounds in this group exhibited varying hydroxylation patterns and were either non-(**6a/b**), mono-(**7a/b**), or dihydroxylated (**9a/b, 10a**). Among the dihydroxylated variants, two structural subtypes were observed: compounds **9a/b** bearing single hydroxyl groups on two distinct aromatic groups and compound **10a**, characterised by a dihydroxyphenyl substituent. The second structural group comprised compounds with an aromatic α-substituent but variation at theβ and γ-positions. Theβ substituent appeared highly flexible and contained either an aliphatic (**8a**/**8b**/**8c**) or indole moiety (**11a**/**11b**/**12a**/**12b**), whilst the γ-substituents were usually aromatic with the exception of an indole at the γ-substituent in **11c**. Compounds **11a/b** differed by a single hydroxyphenyl group proposed to be located on the γ-substituent. Importantly, the position of the substituents at the furanolide core (α-,β, or γ-) is predicted based on the identification of specific ions within MS/MS data (detailed description provided in Supporting Text S3).

**Figure 3:**
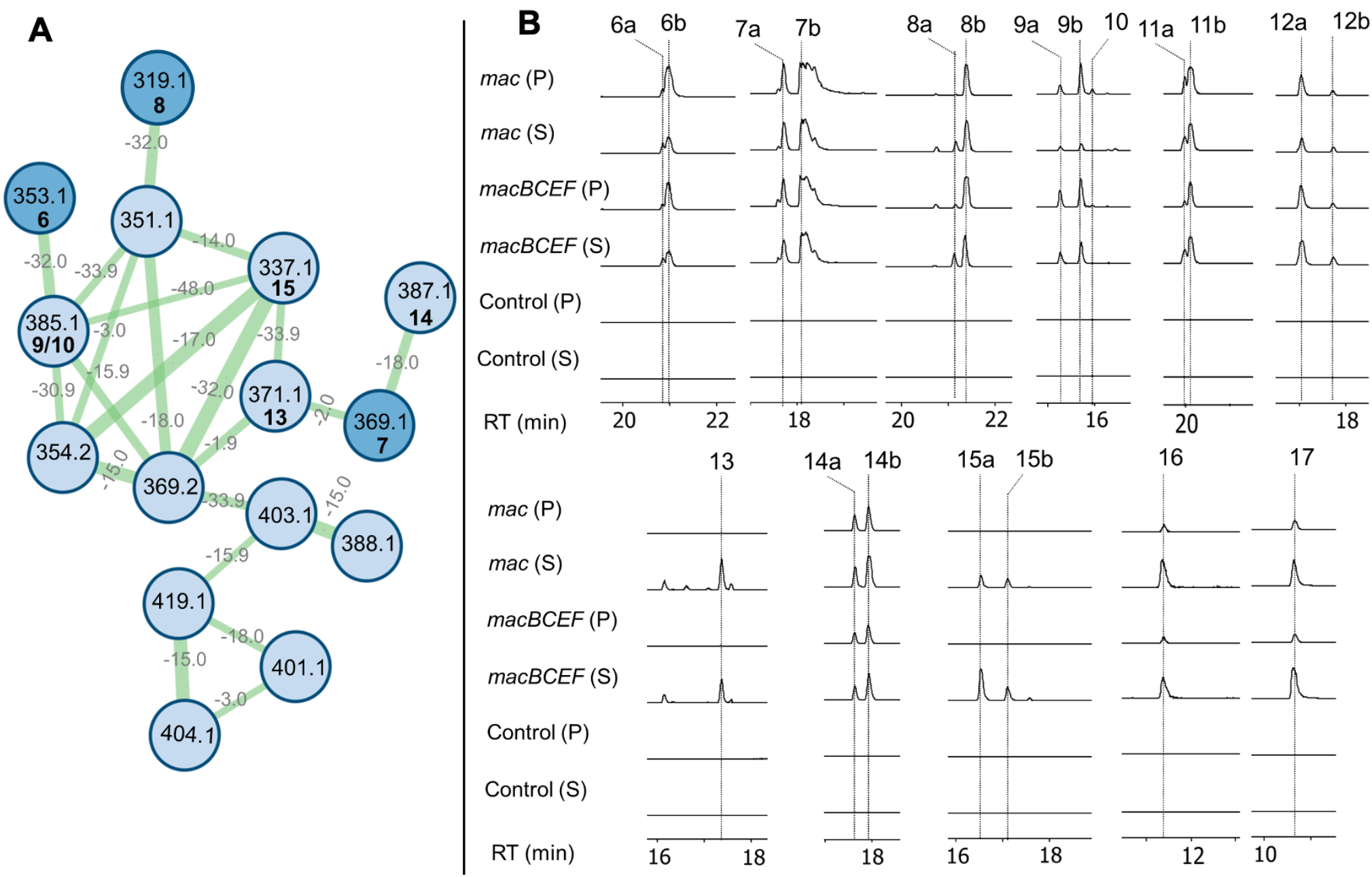
Additional maculalactone analogues and precursors identified by GNPS. **A)** A molecular network of maculalactone NPs created using the online workflow on the GNPS website (http://gnps.ucsd.edu)^30^ and visualised in Cytoscape 3.9.1. Each node represents a parent mass, edges are labelled with the mass difference between metabolites whereas edge thickness is proportional to the cosine similarity score (>0.7). Dark blue nodes indicate compounds **6-8** structurally elucidated by NMR and light blue nodes indicate new compounds proposed to be part of the maculalactone family based on the MS/MS fragmentation patterns. **B)** Extracted ion chromatograms (EICs): **6a/6b** (353.15); **7a/7b** (369.14); **8a/8b, 8c** (319.16); **9a/9b, 10a, 10b** (385.14); **11a/11b, 11c** (392.16); **12a/12b** (408.16); **13a, 13b/13c** (371.16); **14a/14b** (387.16); **15a/15b** (337.18); **16** (257.11); **17** (149.06). All EICs per each panel are depicted in fixed intensity scale. (P) and (S) indicates extracts derived from the pellet and supernatant fractions, respectively. HRMS/MS spectra of all maculalactone-related NPs are fully annotated (Supporting Figures S9-S25).

The third group was characterised by two types of non-aromatic hydroxylations. These modifications, as opposed to hydroxyl substitutions at the aromatic moieties, were implied either by the distinctive loss of H_2_O or by MS/MS fragmentation patterns and key fragment ions compared to previously characterised compounds (Supporting Text S3). Compound **13a** has the same number of carbon and oxygen atoms as **7** but has a 2 Da mass difference. Thus, it is suggested **13a** is hydroxylated on the furanolide ring, leading to saturation of the double bond at position C2-C3 (Supporting Text S3; Supporting Figure S19). Further, an intense fragment ion consistent with the loss of H_2_O supports the idea for tertiary alcohol prone to elimination, in contrast to aromatic hydroxylation. Fragment ions at *m/z* 105.07 and *m/z* 119.05 suggest a saturated single bond between C4 of the furanolide core and the benzylic carbon of the γ-substituent. Notably, **13a** has the same mass as the reported compound maculalactone L, which possesses an α-hydroxylation on the furanolide ring. Other examples of furanolides hydroxylated at the same position are also known, including angiolactone^31^, deoxyenhygrolide I,^32^ and deoxyenhygrolide J^32^. However, based on the MS/MS data of **13a**, the exact location of the hydroxylation cannot be deduced. Secondly, LCMS/MS fragmentation patterns suggested benzylic hydroxylation in some detected analogues (**10b**/**13b**/**13c**/**14a**/**14b**/**15a**/**15b**) (Supporting Text S3). Predicted hydroxylation at the benzyl carbon of either the α-,β, or γ-substituent is supported by the exclusion of saturated furanolide rings consistent with molecular weights and the loss of H_2_O is not consistent with **13a** (tertiary alcohol) but is instead likely a secondary alcohol. Benzylic alcohols have previously been reported for furanolides from myxobacteria, including the deoxyenhygrolide G, H and J.^32^

Maculalactones produced by *K. maculans* structurally fall into two broad categories: the tribenzylbutyrolactones (e.g. maculalactone A; **4**), and the dibenzyldiphenyl-4,5,6,7-tetrahydrobenzofuranones, characterised by a six-membered carbocycle and diverse hydroxylations (e.g. maculalactone F; **5**).^10,12^ However, we were unable to detect any of the latter in our *E. coli* heterologous expression extracts. Furthermore, despite extensive screening, no maculalactone-related NPs could be identified in either the cell pellet or culture supernatant of *Nodularia* sp. NIES-3585. Genetic analysis of the *mac* BGC did not reveal putative genes responsible for the complex dibenzyldiphenyl-4,5,6,7-tetrahydrobenzofuranone scaffold formation and thus, the biosynthesis of these structurally intriguing maculalactones remains unsolved. Based on data obtained in this study, we therefore hypothesise that the *Nodularia* sp. NIES-3585 *mac* BGC is distinct from that encoded in *K. maculans*.

## 3. Conclusion

Furanolides are a class of γ-alkylidenebutenolide NPs with significant structural and functional diversity. Employing the DiPaC strategy, we successfully accessed a cryptic BGC from *Nodularia* sp. NIES-3585 and identified the encoded products in an *E. coli* heterologous host. This work represents the first comprehensive characterisation of the *mac* BGC, providing crucial insights into the genetic basis of maculalactone biosynthesis and expanding our understanding of furanolide diversity in cyanobacteria.

By heterologously expressing the cryptic furanolide BGC, we enabled the discovery and structural elucidation of the new compound maculalactone N (**7**), characterised by a 4-hydroxyphenyl substituent at theβposition. Additionally, we isolated the known compounds maculalactone B (**6**) and deoxyenhygrolide A (**8**), confirming the flexible activity of the *mac* BGC. Whilst these compounds did not exhibit detectable antimicrobial activity under our assay conditions, this does not preclude other biological functions, as the maculalactone family has demonstrated diverse activities including antifouling properties.

The subsequent GNPS analysis expands known maculalactone structural variations, identifying a comprehensive set of 25 maculalactone-related NPs produced by the *mac* BGC. Our study revealed a consistent pattern of aromatic substitution at the α- and γ-positions, whilst demonstrating remarkable flexibility at theβsubstituent, which can accommodate aromatic, indole, or aliphatic substituents. The detection of furan-ring hydroxylated compounds further highlights the structural plasticity of these NPs.

This study not only extends the known chemical space of furanolides but also demonstrates the power of genome mining and heterologous expression techniques to uncover cryptic cyanobacterial biosynthetic potential. Our findings underscore the enzymatic versatility of the maculalactone biosynthetic pathway and provide a foundation for future biocatalytic production of structurally diverse unnatural furanolides through substrate engineering and enzyme promiscuity exploitation.

## 4. Materials and Methods

### 4.1 Bacterial Strains, Plasmids, and Genomic DNA Extraction

*Nodularia* sp. NIES-3585 was purchased from the Japanese Microbial Culture Collection at the National Institute for Environmental Studies (https://mcc.nies.go.jp). The strain was cultured in static batch cultures in BG-11 medium, pH 8 (Sigma-Aldrich, Germany), at room temperature under natural light conditions. Biomass was harvested by centrifugation (4,000*g*, 30 min, 4°C) and frozen at -80°C. Genomic DNA was obtained using an optimised method for cyanobacteria as previously described.^13,14^ Bacterial strains and plasmids used and generated in this study are summarised in Supporting Tables S3 and S4, respectively.

### 4.2 Direct Pathway Cloning

Putative furanolide biosynthetic genes in *Nodularia* sp. NIES-3585 were identified using the *cyb* CybBCEF protein sequences (GenBank accession number BK059219) as a query. Prior to initiating cloning experiments, the intergenic regions of the 15.5 kb *mac* cluster were analysed for potential *E. coli* transcriptional terminators using ARNold,^33^ but no such sequences were identified. Subsequently, primers were designed for targeting the complete predicted *mac* BGC and the minimal *macBCEF* with 20-25 bp homology overlaps at terminal target regions (Supporting Table S5). Attempts to amplify the entire *mac* BGC in one fragment failed, so insert amplification was performed in two fragments (Figure 2). The vector backbone (pET28b-ptetO-GFPv2) was amplified as a linear nucleotide sequence by PCR using previously described primers.^16^ Amplifications were performed in 50 μL reactions using Q5 High-Fidelity DNA polymerase (NEB). Reaction setup and thermocycling conditions were performed according to the manufacturer’s guidelines. For primers containing a homology arm, the best annealing temperatures were evaluated by performing a temperature gradient between 50-65 °C whilst the annealing temperatures for specific primers were calculated using the NEB Tm Calculator (https://tmcalculator.neb.com). The linearised vector was treated with DpnI (37°C, 3 h; 65°C, 20 min) to remove template plasmid DNA.

All DNA fragments were purified directly on column using the Monarch® PCR & DNA Cleanup Kit (NEB), as PCR reactions yielded specific products. For the construction of pET28b-ptetO::*mac* and pET28b-ptetO::*macBCEF*, the purified DNA fragments (0.02-0.5 pmol each) were assembled using either Sequence and Ligation Independent Cloning (SLIC) or HiFi DNA Assembly Cloning Kit (NEB).^14,15^ SLIC assembly was performed in a 10 μL reaction using NEBuffer 2.1 and T4 DNA polymerase (NEB), followed by incubation for 45 s at room temperature and 10 min on ice. HiFi assembly was performed according to the manufacturer’s guidelines, except for the reaction being performed in a 10 µL final volume. For both assembly strategies, half of the DNA assembly reaction (5 µL) was transformed into chemically competent *E. coli* DH5α or *E. coli* DH10 β . Positive clones were identified by colony PCR and sequences verified by restriction digest analysis (ScaI/SspI double digest for pET28b-ptetO::*mac*; DraIII for pET28b-ptetO::*macBCEF*). In addition, the sequences were validated by whole-plasmid sequencing using Oxford Nanopore technology.

### 4.3 Heterologous expression

Each expression experiment was compared to a simultaneous experiment with *E. coli* BAP1 harbouring the pET28b-ptetO-GFPv2 vector backbone, which served as the negative control. A single colony of *E. coli* BAP1 freshly transformed was used to inoculate a 10 mL pre-expression culture grown in LB medium supplemented with 50 µg/mL kanamycin for plasmid selection. The pre-culture was incubated overnight at 30°C with shaking at 200 rpm. The expression cultures for initial verification of recombinant production of new NPs were then inoculated with 1% (v/v) of the pre-expression cultures in 200-400 mL of M9, LB, and TB medium supplemented with 50 µg/mL kanamycin. Incubation was conducted at 37°C with shaking at 200 rpm until an OD_600_ of 0.4 (M9), 0.6 (LB) or 1.0 (TB) was reached. Cultures were then cooled on ice for >1 hour, induced with 0.5 µg/mL tetracycline, and incubated for 5 days at 20°C with shaking at 200 rpm protected from light. Larger expression cultures for compound isolation and characterisation were prepared as above, with 1 L fermentation volume using TB medium for optimal expression.

### 4.4 Extraction and analysis by HPLC

*E. coli* biomass was harvested by centrifugation (10,000*g* for 10 min). The supernatant was transferred to a separating funnel and extracted three times with identical volume of ethyl acetate. The cell biomass was extracted using methanol with simultaneous mechanical cell disruption in a sonicator bath for 30 min. Extracted cell debris was removed by centrifugation (10,000*g* for 10 min). Solvents from both extractions were removed *in vacuo* at 40°C using a rotary evaporator. Desiccated extracts were dissolved in 500 µL HPLC-grade methanol and the resulting solution filtered through a Millex-GP 0.22 µm PES syringe membrane filter (Millipore, USA) prior to injection into HPLC and HPLC-MS systems. *Nodularia* sp. NIES-3585 biomass was collected as a dense culture after one month of growth. Cyanobacterial biomass was centrifuged at 12,000*g* for 30 min to separate the pellet and supernatant. Each fraction was then extracted as described above for *E. coli*.

Analytical HPLC was performed on a Knauer Azura system. Chromatographic separation was performed at 25°C on a Eurosphere II 100-3 C18 A (150 × 4.6 mm) column with integrated precolumn manufactured by Knauer. A wavelength of 220 nm was used for detection and a PDA UV spectrum covering 200-600 nm recorded over the entire run. Eluents used for chromatographic separation were H_2_O (A) and acetonitrile (B), both supplemented with 0.05% trifluoroacetic acid. The gradient was set as follows: Preconditioning 5% B; 0–2 min: 5% B, 2-30 min: 5-100% B, 30-35 min: 100% B, 35-38 min: 100-5% B, at a constant flow rate of 1 mL/min.

### 4.5 Liquid Chromatography-High Resolution Mass Spectrometry (LC-HRMS)

The LC-HRMS experiments were carried out using an Impact II (Bruker Daltonics GmBH & Co. KG) quadrupole time-of-flight (qTOF MS) instrument equipped with an electrospray ionisation source (ESI, Apollo II) coupled to an Elute UHPLC 1300 system (Bruker). An Intensity Solo C18 (1.8 μm, 2.1 mm, 100 mm) from Bruker Daltonics was used for separation. The mobile phases consisted of H_2_O (100%) containing 0.1% formic acid (solvent A) and acetonitrile (100%) containing 0.1% formic acid (solvent B). The following gradient was applied: 0–2 min: 5% B, 2–25 min; 5%–95% B, 25–28 min; 95% B, 28–30 min; 5% B. The flow rate was 0.3 mL/min and the column temperature was set at 40°C. The ESI source was also connected to an external syringe pump (Hamilton syringe 2.5 mL) for pre-acquisition calibration and was operated in the positive ESI mode (ESI+). The external syringe pump was used for mass calibration using a Na-Formate calibrant solution (Na-Formate cluster: 12.5 mL H_2_O, 12.5 mL isopropanol, 50 μL formic acid conc., 250 μL NaOH 1M). The syringe pump injects the calibration solution matching the polarity of ionisation and calibrates the mass axis of the Impact II system in all scan functions used (MS and/or MS/MS). The Q-TOF HRMS method consisted of a full scan TOF survey (50–1300 Da) and a maximum number of three DDA MS/MS scans. The source parameters were as follows: Dry gas 8 L/min, nebulizer gas 1.8 bar, capillary voltage 4.5 kV and end plate voltage of 500 V. For the DDA MS/MS experiments, a collision energy (CE) ramp of 20-50 V was applied. The instrument was controlled by HyStar and Qtof Control software, while data processing and metabolomic analyses were carried out using Data Analysis (Version 6.1) and MetaboScape (Version 8.0.2) software (Bruker).

Principal Component Analysis (PCA) and Partial Least Squares-Discriminant Analysis (PLS-DA) analyses were performed in the MetaboScape software using the following parameters: Feature detection (peak picking) and alignment was done using the T-Rex 3D algorithm provided within MetaboScape with an intensity threshold of 1500 counts and a minimal peak length of 8 data points. For gap filling, the Recursive Feature Extraction algorithm was enabled. Ion deconvolution and de-adducting was performed by incorporating the most common adducts in positive mode (M+H^+^, M+Na^+^, M+K^+^, M-H_2_O+H^+^, M-2H_2_O+H^+^). For computing the final PCA and PLS-DA plots, data were scaled using Pareto scaling.^23,24^

### 4.6 GNPS Molecular Networking

All HRMS data files (.*d* format) were converted to the open data format .*mzML* using the MSConvert software from ProteoWizard.^34^ The converted files were uploaded to the Global Natural Product Social (GNPS) environment, where a classical molecular networking method was generated using the online workflow (http://gnps.ucsd.edu).^35^ MS/MS spectra underwent window-filtering by retaining only the top six fragment ions within the +/-50Da window across the spectrum. The mass tolerance was set to 0.02 Da for both precursor ions and MS/MS fragment ions. A network was constructed with edges filtered according to two criteria: a cosine score threshold above 0.7 and a minimum of five matched peaks. A connection between two nodes was kept only if both nodes ranked among each other’s top 10 most similar matches. Molecular families were limited to a maximum size of 100 nodes, with the lowest-scoring edges removed iteratively until this threshold was met. Networks were then compared against GNPS spectral libraries, with library spectra filtered using identical parameters as for the input data. Network visualization and analysis were performed using Cytoscape 3.9.1.^36^

### 4.7 Isolation of NPs by Semi-Preparative HPLC

Isolation of **6-8** was performed on a semi-preparative HPLC controlled by a Jasco HPLC system consisting of an UV-1575 Intelligent UV/VIS Detector, two PU-2068 Intelligent prep. Pumps, a MIKA 1000 Dynamic Mixing Chamber (1000 μL Portmann Instruments AG Biel-Benken), a LC-NetII/ADC, and a Rheodyne injection valve. The system was controlled by the Galaxie-Software. Chromatographic separation was performed on a Eurosphere II 100-5 C18 A (250 × 16 mm) column with precolumn (30 × 16 mm) provided by Knauer and the eluents used were H_2_O (A) and acetonitrile (B), both supplemented with 0.05% trifluoracetic acid. The separation method was as follows: Preconditioning 5% B; 0-2 min; 5% B, 2-24min; 5-60% B, 24-34min; 60-70% B, 34-35min; 70-100% B,35-40min; 100% B, re-equilibration at 5% B for 7 minutes and at a constant flow rate of 10 mL/min. Fractions were pooled and dried *in vacuo* before NMR analysis.

### 4.7 Determination of Antibacterial Activity

Isolated compounds **6-8** were tested against a range of bacterial strains (Supporting Table S3), using the standard broth two-fold microdilution method in 96-well plates. Each NP was serially diluted in Mueller-Hinton broth to concentrations ranging from 100 to 0.09 μg/mL. An inoculum from overnight cultures from the test strains was adjusted to approximately 5 × 10^5^ CFU/mL and added to each well. The plates were incubated aerobically at 37°C. The minimum inhibitory concentration (MIC) was determined as the lowest concentration of each NP that inhibited microbial growth. Two different protocols were followed: the CLSI M100-S20 guidelines with 16 hours incubation, as previously described^37^, and the EUCAST guidelines (ISO20776-1:2019) with 18–24 hours incubation, according to Ji *et al*.,.^27^ Regular quality control was performed using appropriate reference antibiotics and the experiments were performed in triplicates.

### 4.8 NMR Analysis

NMR spectra were recorded on a Bruker AV-500 spectrometer at 298 K. The chemical shifts are given in δ-values (ppm) and are calibrated on the residual peak of the deuterated solvent (CDCl_3_: δ_H_ = 7.26 ppm, δ_C_ = 77.0 ppm; MeOD-d_4_: δ_H_ = 3.31 ppm, δ_C_ = 49.0 ppm). The coupling constants *J* are given in Hertz [Hz]. The following abbreviations were used for the allocation of signal multiplicities: s – singlet, d – doublet, t – triplet, hept – heptet, m – multiplet.

#### Maculalactone B (6b)

Isolated as a yellow solid. Analytical data in agreement with literature values.^10,12 **1**^**H-NMR** (500 MHz, CDCl_3_): δ = 7.71 (d, *J* = 7.3 Hz, 2 H), 7.35 (t, *J* = 7.5 Hz, 2 H), 7.31– 7.26 (m, 4 H), 7.26–7.23 (m, 3 H), 7.20–7.17 (m, 2 H), 7.12 (d, *J* = 7.0 Hz, 2 H), 5.98 (s, 1 H), 3.93 (s, 2 H), 3.74 (s, 2 H). ^**13**^**C-NMR** (126 MHz, CDCl_3_): δ = 170.5, 150.9, 148.3, 137.6, 136.7, 133.1, 130.6, 129.1, 128.9, 128.8, 128.7, 128.3, 128.0, 127.2, 126.8, 110.6, 30.7, 29.9. **HRMS** (ESI-TOF) *m/z*: [M+H]^+^ calculated for C_25_H_21_O_2_ 353.1542, found 353.1536; [M+Na]^+^ calculated for C_25_H_20_NaO_2_ 375.1361, found 375.1356.

#### Maculalactone N (7b)

Isolated as a yellow solid. ^**1**^**H-NMR** (500 MHz, CDCl_3_): δ = 7.71 (d, *J* = 7.3 Hz, 2 H), 7.35 (t, *J* = 7.5 Hz, 2 H), 7.31–7.26 (m, 2 H), 7.25 (t, *J* = 7.2 Hz, 2 H), 7.20–7.17 (m, 2 H), 6.97 (d, *J* = 8.5 Hz, 2 H), 6.75 (d, *J* = 8.6 Hz, 2 H), 5.98 (s, 1 H), 3.86 (s, 2 H), 3.73 (s, 2 H). ^**13**^**C-NMR** (126 MHz, CDCl_3_): δ = 170.6, 154.7, 151.4, 148.3, 137.6, 133.1, 130.6, 129.5, 129.0, 128.9, 128.82, 128.77, 128.7, 127.7, 126.8, 115.9, 110.7, 29.89, 29.86. **HRMS** (ESI-TOF) *m/z*: [M+H]^+^ calculated for C_25_H_21_O_3_ 369.1491, found 369.1502; [M+Na]^+^ calculated for C_25_H_20_NaO_3_ 391.1310, found 391.1294.

#### Deoxyenhygrolide A (8b)

Isolated as a yellow solid. ^**1**^**H-NMR** (500 MHz, MeOD-d_4_): δ 7.44–7.17 (m, 10H), 6.93 (d, *J* = 1.0 Hz, 1H), 3.76 (s, 1H), 3.71 (s, 2H), 2.32 (d, *J* = 7.2 Hz, 2H), 1.20 (hept, *J* = 6.9 Hz, 1H), 0.46 (d, *J* = 6.7 Hz, 6H). ^**13**^**C-NMR** (126 MHz, MeOD-d_4_): 170.2, 150.6, 149.8, 137.9, 133.9, 132.4, 130.0 (2C), 129.4 (2C), 129.1 (5C), 126.8, 115.8, 35.4, 30.2, 28.7, 21.7. **HRMS** (ESI/QTOF) *m/z*: [M+H]^+^ calcd for C_22_H_23_O_2_^+^ 319.1693, found 319.1676; [M+Na]^+^ calcd for C_22_H_22_NaO_2+_ 341.1512, found 341.1494.

### 4.10 Data Availability

The genome of *Nodularia* sp. NIES-3585 is publicly available in NCBI (accession: BDUB00000000).^38^ The *mac* BGC, pET28b-ptetO::*mac* and pET28b-ptetO::*macBCEF* sequences were submitted to NCBI (accession: BK070015, PV097226 and PV097227, respectively). MS data is available on the MassIVE GNPS repository (accession: MSV000097078).

