## Supporting Information for "The Maculalactone Biosynthetic Gene Cluster, a Cryptic Furanolide Pathway Revealed in *Nodularia* sp. NIES-3585"

**Table S1: Predicted gene function of the maculalactone (*mac*) biosynthetic gene cluster.**

| Locus Tag | Gene | Size (bp) | Predicted Gene Function | Identity/similarity with *cyb* |
|---|---|---|---|---|
| *Ga0262663_114493* | *orf1* | 1167 | DUF7005 family protein | |
| *Ga0262663_114494* | *macA* | 1095 | Deoxy-D-arabinoheptulosonate-7-phosphate-synthase (DHAP Synthase) | *cybA* 78%/89% |
| *Ga0262663_114495* | *macB* | 1824 | Aromatic amino acid-ammonia-lyase | *cybB* 74%/85% |
| *Ga0262663_114496* | *macC* | 1539 | Long-chain acyl-CoA-synthetase | *cybC* 71%/85% |
| *Ga0262663_114497* | *macE* | 1812 | Thiamine pyrophosphate-binding protein (TPP) | *cybE* 65%/78% |
| *Ga0262663_114498* | *macF* | 1134 | Furanolide synthase | *cybF* 76%/88% |
| *Ga0262663_114499* | *macM* | 417 | Chorismate mutase | |
| *Ga0262663_114500* | *macK* | 1320 | Tyrosinase | *cybK* 47%/64% |
| *Ga0262663_114501* | *orf2* | 942 | Hypothetic protein | |
| *Ga0262663_114502* | *orf3* | 630 | Putative NADH-flavin reductase | |
| *Ga0262663_114503* | *orf4* | 738 | Phycocyanobilin:ferredoxin oxidoreductase | |

**Table S2: $^{1}$H and $^{13}$C NMR data for 6, 7 and 8.** Compounds **6** and **7** were measured in $CDCl_3$. Compound **8** was measured in MeOD-$d_4$. Abbreviations: s: singlet, d: doublet, t: triplet, hept: heptet, m: multiplet, *13C signals extracted from 2D NMR data.

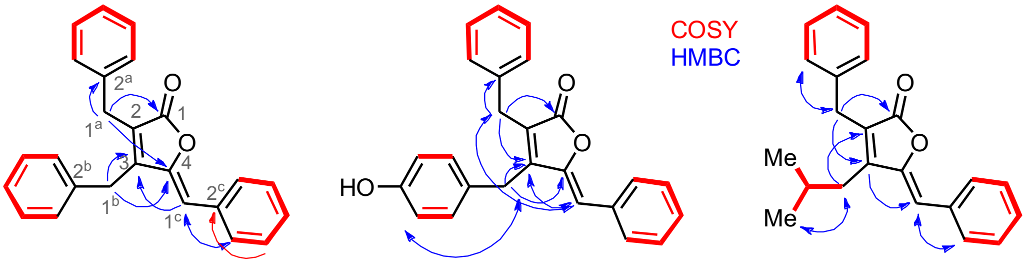

6 7 8

| Atom | δC (ppm) 6 | 7 | 8* | δH (multiplicity, coupling constants in Hz) 6 | 7 | 8 |
|---|---|---|---|---|---|---|
| 1 | 170.5 | 170.6 | 170.2 | | | |
| 2 | 128.0 | 128.8 | 132.4 | | | |
| 3 | 150.9 | 151.4 | 150.6 | | | |
| 4 | 148.3 | 148.3 | 149.8 | | | |
| 1ª | 29.9 | 29.9 | 30.2 | 3.74 (s) | 3.73 (s) | 3.71 (s) |
| 2ª | 137.6 | 137.6 | 137.9 | | | |
| 3ª/3ª' | 128.7 | 128.7 | 129.1 | 7.20–7.17 (m) | 7.20-7.17 (m) | 7.44–7.32 (m) |
| 4ª/4ª' | 128.8 | 128.9 | 129.4 | 7.31–7.26 (m) | 7.31-7.26 (m) | 7.44–7.17 (m) |
| 5ª | 126.8 | 126.8 | 126.8 | 7.26–7.23 (m) | 7.25 (t, 7.2) | 7.44–7.17 (m) |
| 1ᵇ | 30.7 | 29.9 | 35.4 | 3.93 (s) | 3.86 (s) | 2.32 (d, 7.2) |
| 2ᵇ | 136.7 | 129.0 | 28.7 | | | 1.20 (hept, 6.9) |
| 3ᵇ/3ᵇ' | 128.3 | 129.5 | 21.7 | 7.12 (d, 7.0) | 6.97 (d, 8.5) | 0.46 (d, 6.7) |
| 4ᵇ/4ᵇ' | 129.1 | 115.9 | | 7.31–7.26 (m) | 6.75 (d, 8.6) | |
| 5ᵇ | 127.2 | 154.7 | | 7.26–7.23 (m) | | |
| 1ᶜ | 110.6 | 110.7 | 115.8 | 5.98 (s) | 5.98 (s) | 6.93 (s) |
| 2ᶜ | 133.1 | 133.1 | 133.9 | | | |
| 3ᶜ/3ᶜ' | 130.6 | 130.6 | 130.0 | 7.71 (d, 7.3) | 7.71 (d, 7.3) | 7.44–7.32 (m) |
| 4ᶜ/4ᶜ' | 128.8 | 128.8 | 129.1 | 7.35 (t, 7.5) | 7.35 (t, 7.5) | 7.44–7.17 (m) |
| 5ᶜ | 128.9 | 127.7 | 129.1 | 7.26–7.23 (m) | 7.25 (t, 7.2) | 7.44–7.17 (m) |

**Table S3. Bacterial strains used in this study.**

| Strains | Description | Reference or Source‡ |
|---|---|---|
| *Nodularia* sp. NIES-3585 | Native producer of maculalactones. | NIES |
| *Escherichia coli* DH5α | Host strain for cloning. | NEB GmbH |
| *Escherichia coli* DH10β | Host strain for cloning. | Durfee *et al.,* (2008)[1] |
| *Escherichia coli* BAP1 | Heterologous expression host and reference strains for antibacterial susceptibility testing performed following CLSI guidelines. | Pfeifer *et al.,* (2001)[2] |
| *Escherichia coli* CCM2024 | Reference strains for antibacterial susceptibility testing performed following CLSI guidelines. | CCM |
| *Bacillus subtilis* CCM1999 | | CCM |
| *Pseudomonas aeruginosa* CCM1959 | | CCM |
| *Staphylococcus aureus* CCM3824 | | CCM |
| *Streptococcus sanguinis* CCM4047 | | CCM |
| *Escherichia coli* BW25113 | Reference strains for antibacterial susceptibility testing performed following EUCAST guidelines. | ECGRC |
| *Escherichia coli* Δ*tolC* | | HIPS-MINS |
| *Escherichia coli* DSM-1116 | | DSMZ |
| *Klebsiella pneumoniae* DSM-30104 | | DSMZ |
| *Pseudomonas aeruginosa* DSM-19882 (PA14) | | DSMZ |
| *Pseudomonas aeruginosa* Δ*mexAB* (PA14) | | HIPS-MINS |

‡NIES - National Institute of Environmental Studies Culture Collection.
‡DSMZ - German Collection of Microorganisms and Cell Cultures.
‡CCM - Czech Collection of Microorganisms.
‡ECGRC - *E. coli* Genomics Resource Centre.
‡HIPS-MINS - Helmholtz Institute for Pharmaceutical Research Saarland (HIPS), Department of Microbial Natural Products (MINS).

**Table S4. Plasmids used in this study.**

| Plasmids | Description | Reference or Source |
|---|---|---|
| pET28b-ptetO-GFPv2 (6,029 bp) | Tetracycline inducible expression plasmid, ColE1, $Kan^R$, GFP. | Duell *et al.,* (2019)[3] |
| pET28b-ptetO::*mac* (22,090 bp) | Plasmid pET28b-ptetO-GFPv2 harbouring a 15,538 bp insert corresponding to the complete predicted *mac* BGC. GFP is located downstream of the insert. | This study. Accession: PV097226 |
| pET28b-ptetO::*macBCEF* (13,046 bp) | Plasmid pET28b-ptetO-GFPv2 harbouring a 6,493 bp insert corresponding to the core *macBCEF* genes. GFP is located downstream of the insert. | This study. Accession: PV097227 |

**Table S5: List of Primers.**

| Oligonucleotide | Sequence (5' → 3')¥ | Description |
|---|---|---|
| **spec-ptet-R** | GGTCGATCCTCTTCTCTATC | Specific amplification of pET28b-ptetO-GFPv2 vector backbone.[3] |
| **C-GFP_for_1** | CATGGTTAGCAAAGGTGAAG | Specific amplification of pET28b-ptetO-GFPv2 vector backbone.[3] |
| **gib-mac_F** | tcagtgatagagaagaggatcgaccATGGAAAAACGGCAAACATTTC | Initially designed to amplify the full *mac* BGC (15.5 kb). Then used to amplify the upstream fragment (7.5 kb) to generate pET28b-ptetO::*mac.* |
| **gib-mac-frag1_R** | CACAAAAGGAACGGGTGAAG | Designed to amplify the upstream fragment (7.5 kb) of the *mac* BGC to generate pET28b-ptetO::*mac.* |
| **gib-mac-frag2_F** | AGCGTTAGAAAAAGCAATGG | Designed to amplify the downstream fragment (8 kb) of the *mac* BGC to generate pET28b-ptetO::*mac.* |
| **gib-mac_R** | cagttcttcacctttgctaaccatgCTAGCAAAGACTAGGAATTTTTTC | Initially designed to amplify the full *mac* BGC (15.5 kb). Then used to amplify the downstream fragment (8 kb) to generate pET28b-ptetO::*mac.* |
| **gib-macBCEF_F** | tcagtgatagagaagaggatcgaccCATGGGTACACAAGCATTTC | Designed to amplify the minimal furanolide core forming genes *macBCEF* for pET28b-ptetO::*macBCEF* construction. |
| **gib-macBCEF_R** | cagttcttcacctttgctaaccatgCTAGCAAAGACTAGGAATTTTTTC | Designed to amplify the minimal furanolide core forming genes *macBCEF* for pET28b-ptetO::*macBCEF* construction. |
| **screen_ptetF2** | TCCGACCTCATTAAGCAGC | Colony screening primer targeting the ptetO promoter sequence in pET28b-ptetO-GFPv2 vector backbone.[3] |
| **screen_GFP_R** | TTACCGTTGGTCGCATCACC | Colony screening primer targeting the *gfp* sequence in pET28b-ptetO-GFPv2 vector backbone.[3] |
| **screen-mac_F** | TTTGCATCTTTGTGCGTCCTG | Colony screening primer targeting the *orf4* gene in *mac* sequence. |

**¥Underlined lower-case sequences indicate homology arms.**

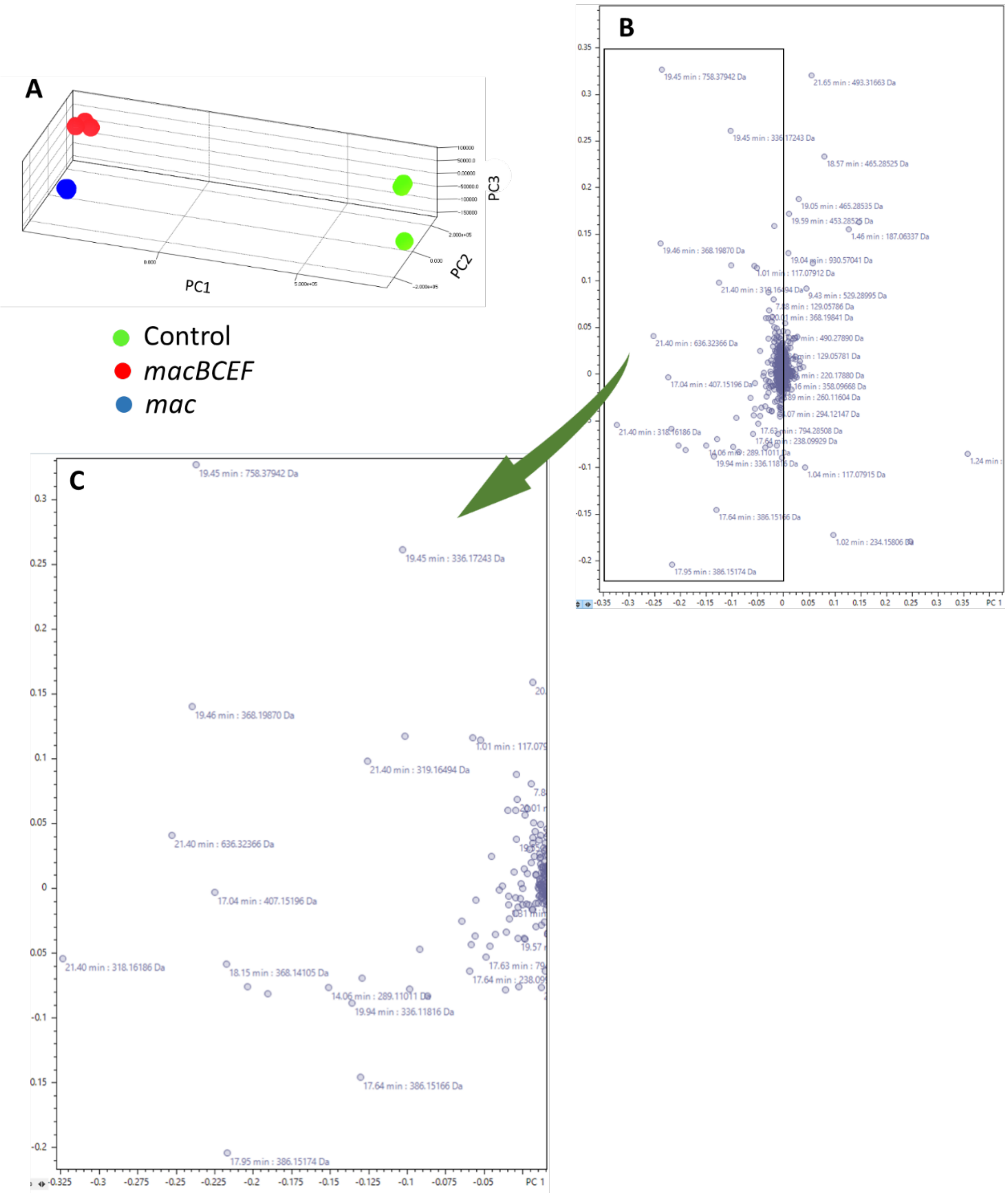


**Figure S1. Principal Component Analysis (PCA) of heterologously expressed constructs in *E. coli*. A)** 3D score plot showing distinct clustering between control (pET28b-ptetO-GFPv2 vector), pET28b-ptetO::*mac* and pET28b-ptetO::*macBCEF.* **B)** Loading plot displaying the distribution of all the metabolites in the data matrix. **C)** Detailed view of the loading plot showing the key metabolites that contribute to the separation between three sample groups (control, *macBCEF*, and *mac*). Cross-validation tests: $R^2$ = 0.879, $Q^2$ = 0.55.

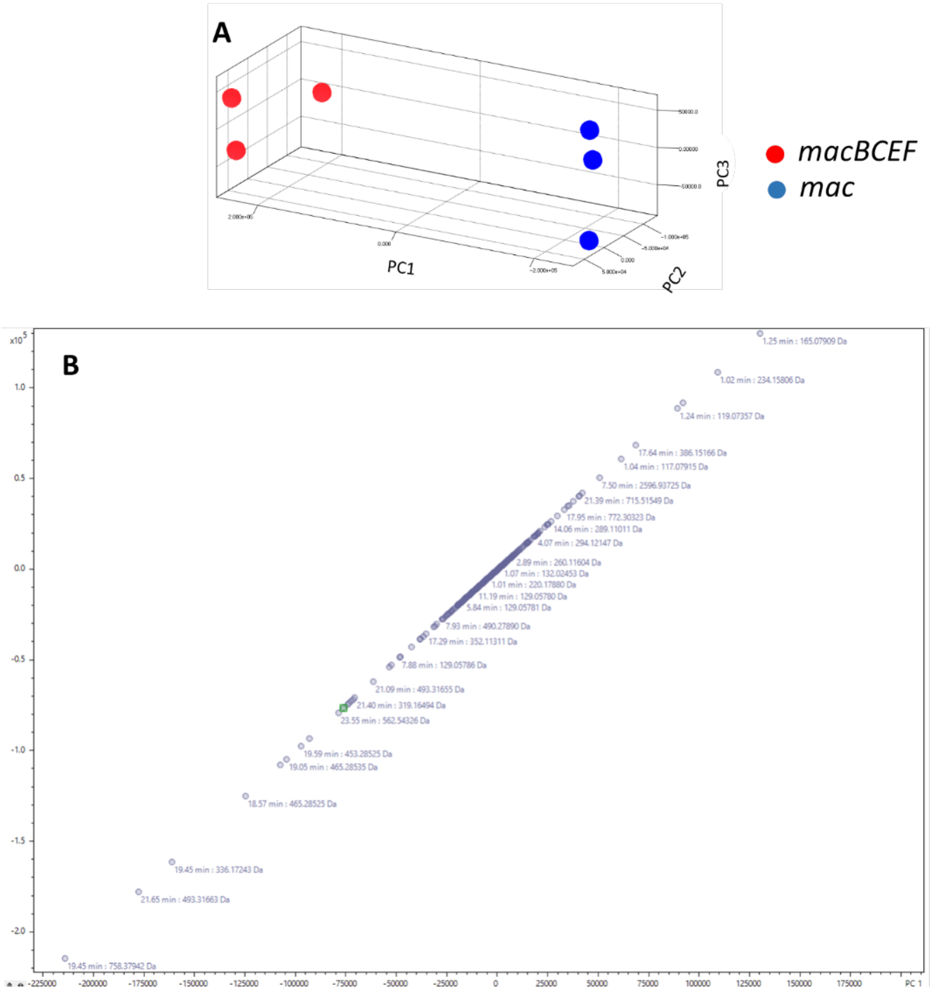


**Figure S2: Partial Least Squares-Discriminant Analysis (PLS-DA) heterologously expressed constructs in *E. coli*. A)** 3D score plot showing distinct clustering between pET28b-ptetO::*mac* and pET28b-ptetO::*macBCEF* transformants. **B)** Loading plot displaying the distribution of all the metabolites in the data matrix. Cross-validation tests: $R^2$ = 0.88, $Q^2$ = 0.43. Permutation test result (for 1000 permutations): p = 0.035.

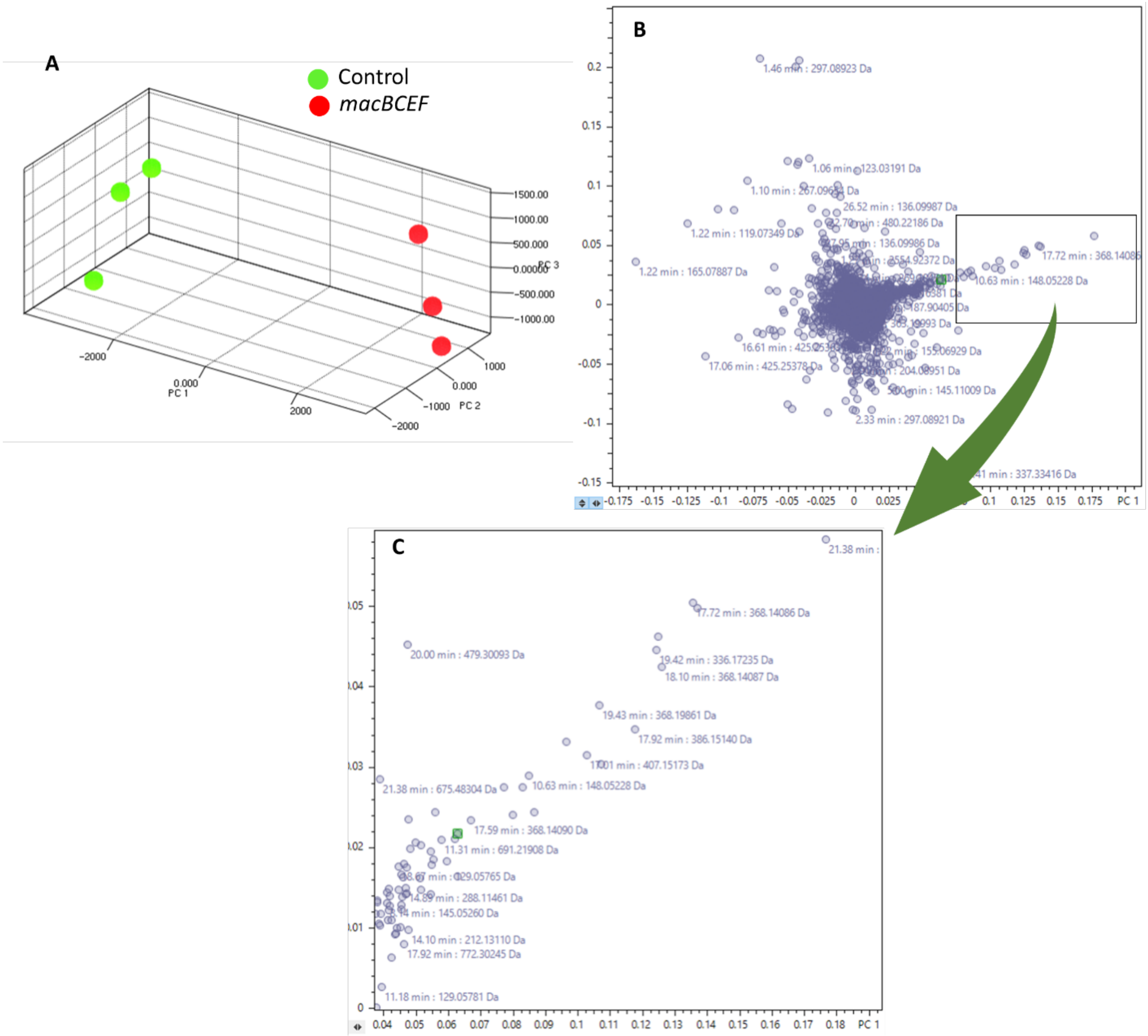


**Figure S3. Principal Component Analysis (PCA) of maculalactone samples: A)** 3D score plot showing distintic clustering between control (pET28b-ptetO-GFPv2 vector) and pET28b-ptetO::*macBCEF*. **B)** Loading plot displaying the distribution of all the metabolites in the data matrix. **C)** Detailed view of the loading plot showing the key metabolites that contribute to the separation between three sample groups (control and *macBCEF*). Cross-validation tests: $R^2$ = 0.97, $Q^2$ = 0.57.

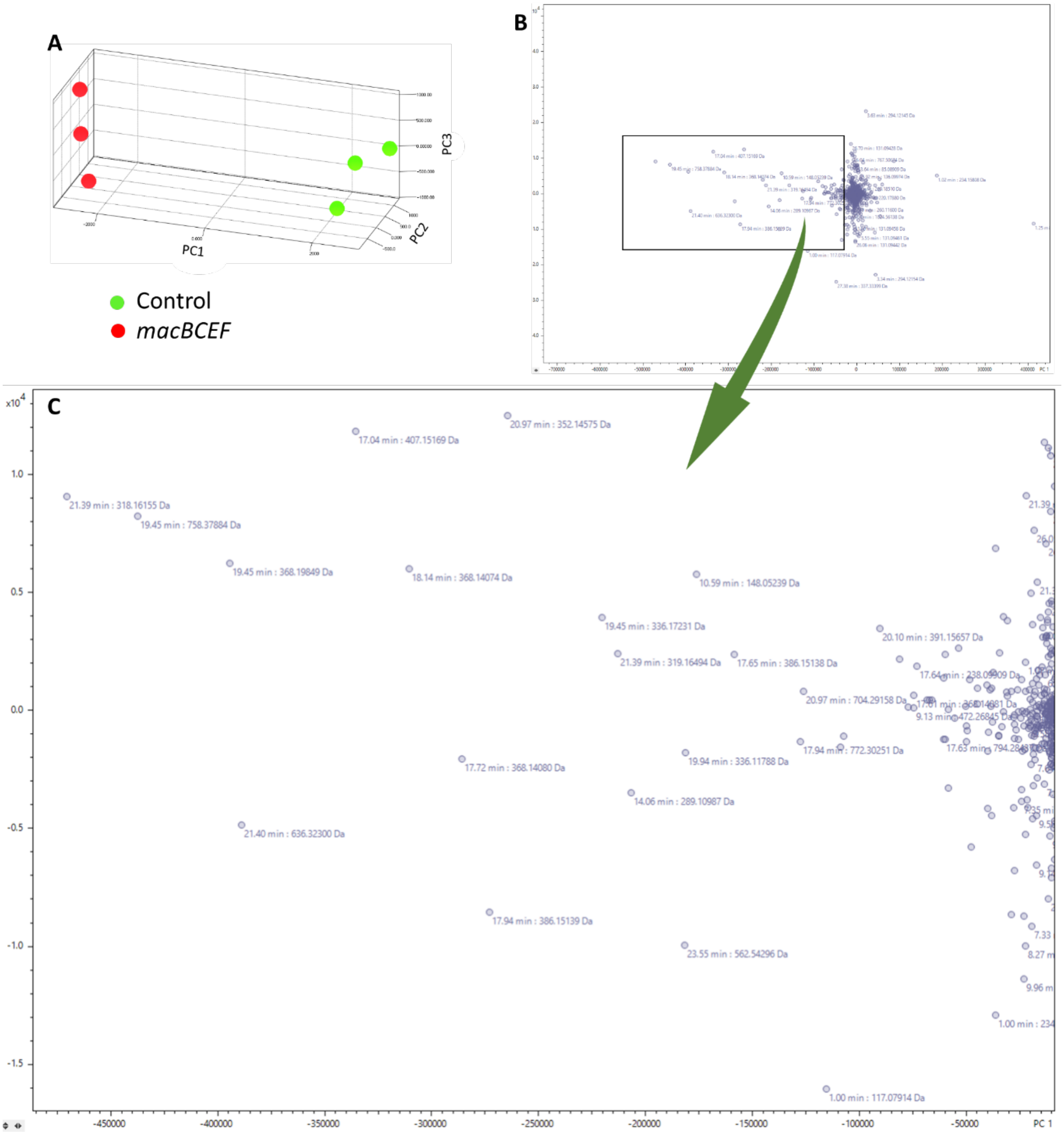


**Figure S4. Partial Least Squares-Discriminant Analysis (PLS-DA) of maculalactone samples:** **A)** 3D Score plot showing clear separation between control (empty vector) and pET28b-ptetO::*macBCEF* transformants **B)** Loading plot showing all the metabolites comprising the whole data matrix, **C)** Zoomed region of loading plot depicting metabolites which contribute the most to the separation between two groups (Control vs. *macBCEF*) of samples. Cross-validation tests: $R^2$ = 0.89, $Q^2$ = 0.47. Permutation test result (for 1000 permutations): p = 0.02.

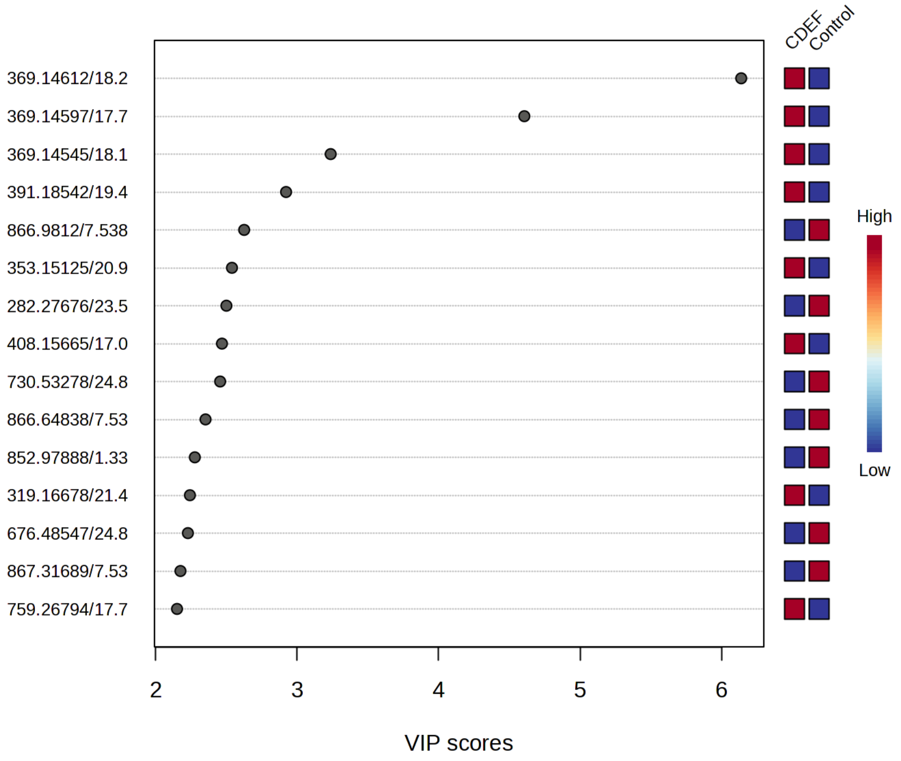


**Figure S5. Variable Importance Projections (VIP) plot.** depicting top contributing metabolites in the group separation between Control and macCDEF inferred from PLS discriminant analysis. Most of the metabolites upregulated in macCDEF (*m*/*z* 369.14; 353.14; 408.15; 319.16, etc) belong to the maculalactone family of furanolides.

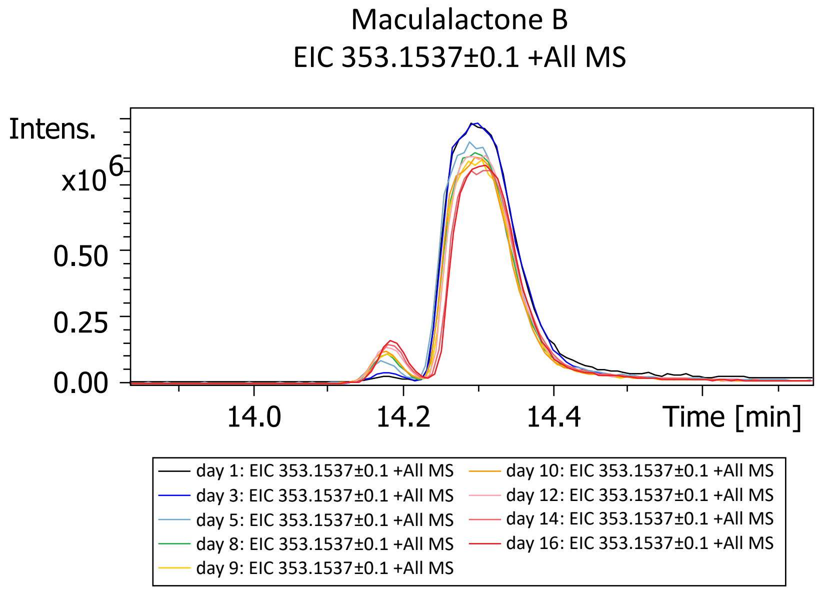


**Figure S6: Interconversion of maculalactone B and maculalactone C *E*/*Z*-isomers.** The major peak of the more thermodynamically stable *Z*-isomer maculalactone B was isolated. Over time there was a gradual conversion to the *E*-isomer maculalactone C.

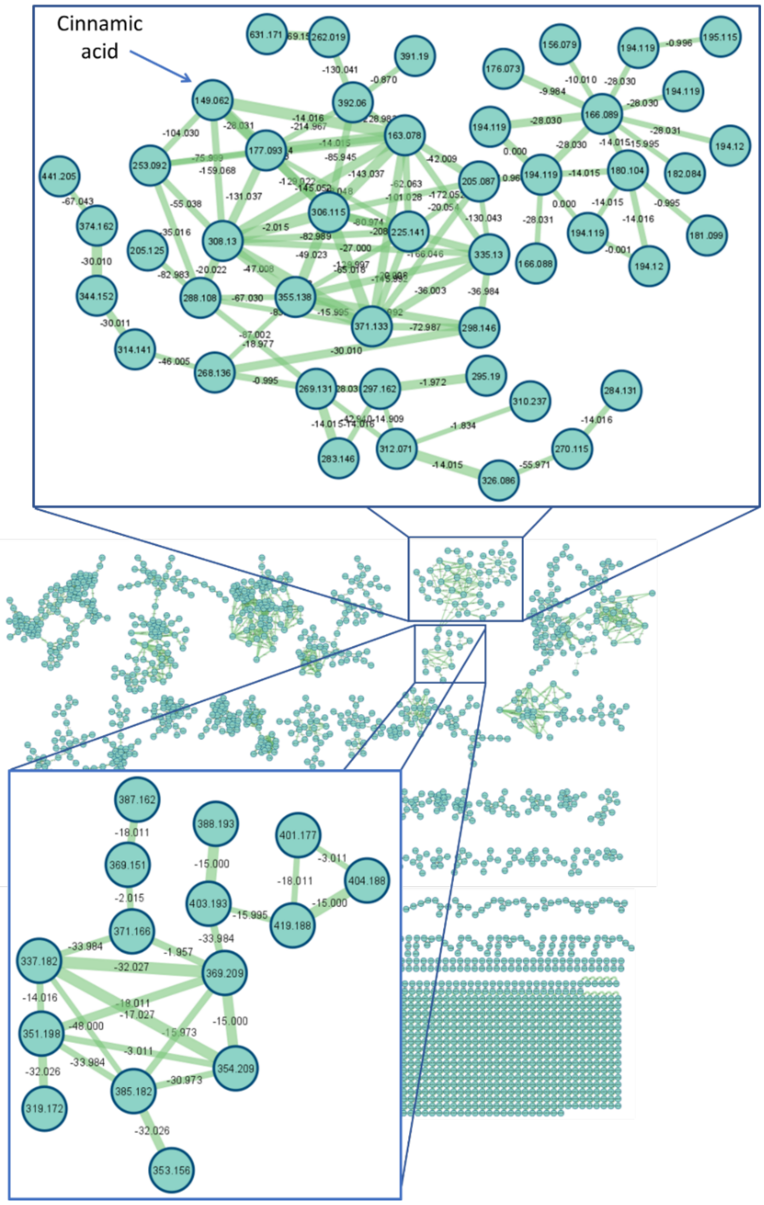


**Figure S7. Full Molecular Network of *E. coli* expressing the maculalactone BGC.** A molecular network was created using the online workflow (https://ccms-ucsd.github.io/GNPSDocumentation/) on the GNPS website (http://gnps.ucsd.edu).[4] The two boxes indicate network harbouring cinnamic acid (top) and compound **6, 7** and **8** (bottom).

#### Supplementary Text S1: MS/MS fragmentation rationale for furanolide natural products exemplified by anhydrocyanobacterin

Anhydrocyanobacterin is a suitable example for this purpose since it contains three different substituents in $\alpha$-, $\beta$-, and $\gamma$-positions, which allow us to predict the sequence of losses of the three substituents as well as other relevant fragments formed in conjunction with individual substituents with the furanolide core. As can be seen, from anhydrocyanobacterin spectrum, the $\alpha$-substituent is cleaved first (-156 Da loss, -$C_7H_5ClO_2$, ring A), which represents a very important step regarding structural elucidation of furanolides and the mass of the $\alpha$-substituent. This step appears common across all furanolides. The largest fragment ion observed (*m*/*z* 257.11) is composed of the furanolide core and both $\beta$- and $\gamma$-substituents. From this point, the ion at *m*/*z* 257.11 undergoes two different paths, decomposition of the furanolide core and/or the parallel cleavage of $\beta$- and $\gamma$-substituents. Decomposition of the furanolide core is documented by two successive losses, 28 Da (keto group as CO) and 18 Da (O atom as water). On the other hand, loss of the isopropyl group in $\beta$-position generates the fragment ion at *m*/*z* 227.07, whereas cleavage of the methoxyphenyl group (ring C) affords fragment ion at *m*/*z* 149.06. An important diagnostic ion seems to be fragment at *m*/*z* 187.07 which is composed of the $\beta$-substituent (isopropyl group), $\gamma$-substituent (ring C) as well as the connecting carbons (C3-C4 of the furanolide core and the benzylic carbon) between these two moieties. Fragments with *m*/*z* 121.06 and *m*/*z* 135.04 are also important ions since both give information regarding $\gamma$-substituent (ring C). These are key fragments in anhydrocyanobacterin fragmentation which uncover the canonical gas phase behaviour of furanolide natural products. This fragmentation rationale exemplified by anhydrocyanobacterin is supported and observed in multiple other examples of MS/MS spectra of cyanobacterin- and maculalactone-like furanolide derivatives produced chemo-enzymatically within our laboratory and published elsewhere.[5–7]

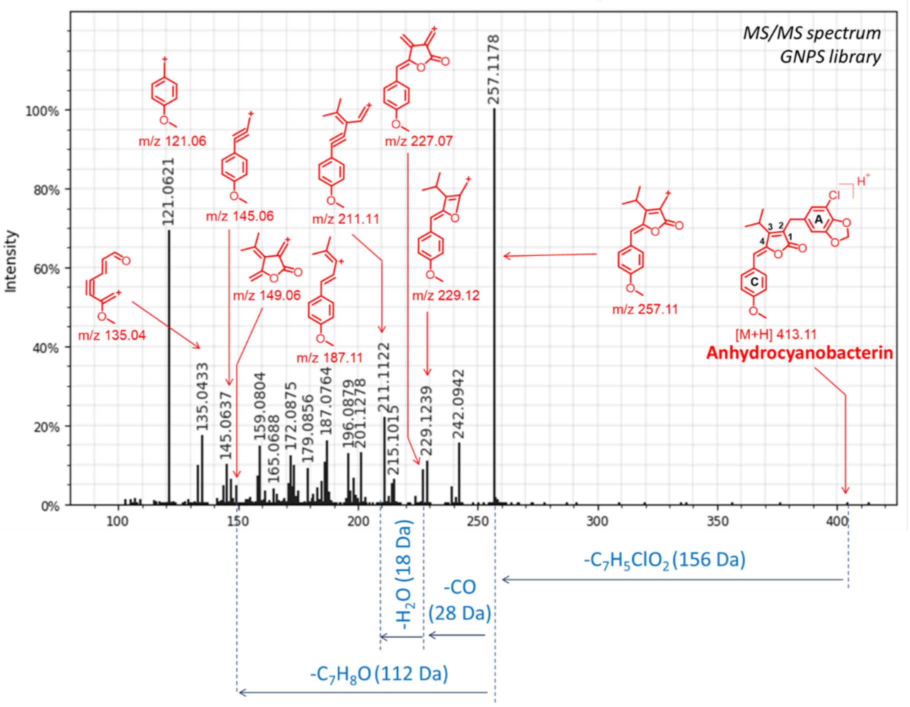


**Figure S8. HRMS/MS spectrum of the known furanolide metabolite, anhydrocyanobacterin and its fragmentation model.** This spectrum was downloaded from the GNPS library and the image was generated using USI Resolution tool within GNPS environment.

**Supplementary Text S2: MS/MS fragmentation rationale applied to compounds maculalactone B/C (6), maculalactone N (7) and deoxyenhygrolide A/B (8)**

Based on the fragmentation patterns inferred from anhydrocyanobacterin (Supporting Text S1), the fragmentation of rationale for the isolated compounds **6**-**8** was built (Supporting Figure S9-11). Compound **6** ($C_{25}H_{21}O_2^+$) follows the same fragmentation logic as described for anhydrocyanobacterin (Figure S8). Despite **6** containing three identical substituents and therefore cleavage of the respective substituents will generate ions with identical mass, it is known that the α-substituent will be lost first affording the fragment ion at *m*/*z* 275.10. In addition to the furanolide core decomposition, parallel cleavages of the β- (ring B) and γ-substituent (ring C) from the fragment ion at *m*/*z* 275.10 was also observed. Noticeably, both these cleavages generate fragments with the same mass *m*/*z* 197.05 which explains the double abundance intensity of *m*/*z* 197.05 compared to the precursor ion (*m*/*z* 353.15). The MS/MS spectrum of **6** also contains the fragment ion at *m*/*z* 105.03, homologous to the fragment ion *m*/*z* 135.04 for anhydrocyanobacterin, indicating a non-substituted γ-substituent (ring C) in the case of **6**. Maculalactone C (**6a**) is the *E*-isomer of **6b** and both exhibit identical MS/MS fragmentation (Supporting Figure S9).

Maculalactone N (**7**, $C_{25}H_{21}O_3^+$) contains one oxygen atom more than **6**. Occurrence of the fragment ion at *m*/*z* 107.05 in the MS/MS spectrum of **7**, representing a hydroxyphenyl motif, provides enough evidence that this oxygen atom (as OH group) is located on one of the aromatic rings (α, β, or γ) of compound **7** (Supporting Figure S10). The initial 78 Da loss (α-substituent) in the MS/MS spectrum of **7**, generating fragment ion at *m*/*z* 291.10 ($C_{19}H_{15}O_3^+$) demonstrates that the hydroxyphenyl group is not located on the α-substituent. Presence of the fragment ion *m*/*z* 105.03, indicates that the γ-substituent is intact and non-substituted. Hence, this supports the location of a hydroxyphenyl located at the β-substituent, which is also supported by the occurrence of other fragment ions such as *m*/*z* 213.05 and *m*/*z* 145.06 and by NMR (Table S2).

Deoxyenhygrolide A/B (**8b**/**8a**, $C_{22}H_{23}O_2^+$) follow an analogical mode of fragmentation as for the abovementioned compounds (Supporting Figure S11). Possessing 3 carbons less compared to **6** indicates a four-carbon alkyl group in its structure. Typical initial 78 Da loss (α-substituent) and the presence of the diagnostic ion at *m*/*z* 105.03 (γ-substituent) suggests that the isobutyl group is connected at the β-position of the furanolide core. NMR analysis confirms an isobutyl moiety incorporated in the chemical scaffold of **8a**/**8b**. Compound **8c** exerts a congruent mode of fragmentation as **8a**/**8b**, with a four-carbon alkyl group in β-position. However, in the case of **8c**, presence of the fragment ion at *m*/*z* 57.07 suggests a *sec*-butyl group instead of an isobutyl group (Supporting Figure S12). The MS/MS differentiation between these two alkyl groups was demonstrated previously, such is the case of discrimination between amino acids leucine and isoleucine.[8]

**Supplementary Text S3: Putative maculalactone-like furanolides based on MS/MS fragmentation rationale**

A list of masses related to compound **6**-**8** were identified using the GNPS molecular network and were analysed within the framework of the newly developed MS/MS-based fragmentation rationale (Table 1; Supporting Figure S12-25). Compound **9** ($C_{25}H_{21}O_4^+$) contains two oxygen atoms more than **6**. Initial 78 Da loss in the MS/MS spectrum of **9** indicates a non-substituted phenyl substituent at the α-position. The presence of *m*/*z* 307 and characteristic ion at *m*/*z* 107.05 in addition to the absence of *m*/*z* 105.03 (indicative of non-substituted phenyl at the γ-position) all suggest the oxygen atoms of compound **9** exist as two hydroxyphenyl functionalities at the β- (ring B) and γ- (ring C) positions. Moreover, the 94 Da opposed to 78 Da loss of ion *m*/*z* 307 suggests the oxygen is lost ($C_6H_6O$) at either the ring B or C. Compound **10a** ($C_{25}H_{21}O_4^+$) possess the same chemical composition as **9**, but with a different MS/MS spectrum. Firstly, initial loss of 78 Da loss (non-substituted phenyl at the α-position), absence of the mono-hydroxylated aromatic (*m*/*z* 107.05) as well as the presence of non-substituted phenyl at the γ-position (*m*/*z* 105.03; ring C) suggests dihydroxylation of the β-substituent. In addition, presence of fragment ions at *m*/*z* 123.04, *m*/*z* 229.05 and *m*/*z* 197.05 all suggest both oxygen atoms are located on the same β-substituent aromatic ring in compound **10a**.

Compound **11a/11b** ($C_{27}H_{22}NO_2^+$) contains two carbons and one nitrogen more than **6**. This chemical composition as well as the presence of the typical 2-methyl-indole cation (*m*/*z* 130.06) indicate an indole group instead of a common aromatic group linked to the furanolide core. Initial 78 Da loss and the *m*/*z* 105.03 fragment ion indicates a non-substituted phenyl rings at the α- and γ-positions, thus suggesting the indole group as the β-substituent. Moreover, occurrence of the ion at *m*/*z* 236.07 as homologous to ion *m*/*z* 213.05 of **7** also improves confidence of the assignment. Compound **11c** possess the same chemical formula as **11a/11b** but a different fragmentation pattern. An indole is again believed to be part of the molecule due to the respective ion *m/z* 130.06. Initial loss of 78 Da, absence of the ion *m*/*z* 105.03, as well as the 117 Da loss of the indole ($C_8H_7N$) motif, clearly suggests the indole group is located at the γ-position. Compounds **12a/12b** ($C_{27}H_{22}NO_3^+$) contain one oxygen atom more than compounds **11a/11b**. Their fragmentation patterns demonstrate that **12** is a mono-hydroxylated analogue of **11a/11b** where the hydroxy group is located on the aromatic at the γ-position. Several fragmentation events support this assignment, such as the non-substituted 78 Da loss (α-substituent), absence of the fragment ion at *m*/*z* 105.03, and presence of the fragment ions at *m*/*z* 107.05 and *m*/*z* 213.05.

*Compounds with alternative hydroxylation patterns*

The strong fragmentation identities of substituents connected to the furanolide core and a subset of hydroxylated analogues possessing the same exact mass, but distinct fragmentation patterns compared to previously identified compounds, were identified. Compound **13a** ($C_{25}H_{23}O_3^+$) contains 2 Da (two protons) more than compound **7**. Moreover, compound **13a** and compound **7**contains one oxygen atom (as OH group) more than **6**. While the hydroxyl group in **7** is connected to an aromatic ring, the 2 Da mass difference in the case of **13a**, suggests that the hydroxyl group is instead connected to the furanolide core, ultimately saturating one of the original double bonds in the lactone ring. In addition, very low abundance of the **13a** precursor ion *m*/*z* 371.16 supports the idea for an aliphatic tertiary alcohol, since tertiary alcohols are very unstable in gas phase and the resulting ion (*m*/*z* 353.15) after water loss is very high compared to the parent ion. Moreover, fragment ions at *m*/*z* 105.07 and *m*/*z* 119.05 suggest a saturated single bond between C4 of the furanolide core and the benzylic carbon of the γ-substituent (ring C).

Compound **10b** ($C_{25}H_{21}O_4^+$), possess the same chemical formula as **9** and **10a** but possesses a different fragmentation pattern. Presence of fragment ion at *m*/*z* 107.05 and *m*/*z* 105.03 and 78 Da loss suggests a hydroxyphenyl and two non-substituted aromatics at the β-, γ- and α-positions, respectively. Using this logic, only a single oxygen has been accounted for. The MS/MS spectrum of **10b** also indicates the successive losses of two water molecules (2 x 18 Da). From the general gas phase behaviour of furanolides, one water molecule is attributed to the decomposition of the furanolide core while the second water molecule suggests a hydroxyl group attached to an benzylic carbon surrounding the furanolide core. A direct connection of this hydroxyl group to the furanolide core ($\alpha$- $\beta$- or $\gamma$-position such as compound **13a**) is excluded, since these bonds would automatically imply a 2 Da increase in the overall mass of the precursor ion and the strong fragmentation from a tertiary alcohol, such as the case with **13a**, is also not observed. With this in mind, the second hydroxyl group of **10b** thought to be a secondary alcohol located on one of the benzylic carbons connected at the $\alpha$- or $\beta$-position. Since, we observe the typical initial 78 Da loss, the position of this hydroxy group is possibly located on the benzylic carbon at the $\beta$-position.

Compounds **13b**/**13c** possess identical chemical composition as **13a**, however a quite different fragmentation mode. Here again we have an aliphatic hydroxylation incorporated in the structure of **13b**/**13c**. An interesting fragment appears at *m*/*z* 221.09 ($C_{16}H_{13}O^+$) homologous to fragment ion *m*/*z* 187.07 in the MS/MS spectrum of anhydrocyanobacterin. The fact that this oxygen atom (as OH group) can be retained in some of the downstream fragments such as the fragment ion *m*/*z* 221.09 suggests it is not a tertiary alcohol (such as compound **13a**), thus most likely represents a secondary alcohol on a benzylic carbon.

Compound **14** ($C_{25}H_{23}O_4^+$) contain 2 Da more than compound **9**. Presence of the hydroxyphenyl cation (*m*/*z* 107.05) in the MS/MS spectra of **14** indicates one of the hydroxy groups is connected to a phenyl ring  According to the previously discussed logic, the secondary hydroxyl group is not a tertiary alcohol and therefore is located on one of the benzylic carbons. The same fragmentation logic as seen in the case of **13b**/**13c** and **14**, can be noticed in the MS/MS spectra of compound **15**. Compound **15** appears to be benzylic hydroxylated analogues of deoxyenhygrolide A/B (**8b**/**8a**), respectively. The strongest evidence for structural assignment of **15** is presence of the fragment ion at *m*/*z* 187.11 in their corresponding spectra, as homologous ions to the fragment ion *m*/*z* 221.09 and *m*/*z* 237.09 of compounds **13b/13c and 14**, respectively. Benzyl alcohols have previously been reported for other furanolides from myxobacteria including the deoxyenhygrolide G, H and J.[9] However, a biosynthetic basis for their presence was never investigated and so far hydroxylation at this position remains elusive.

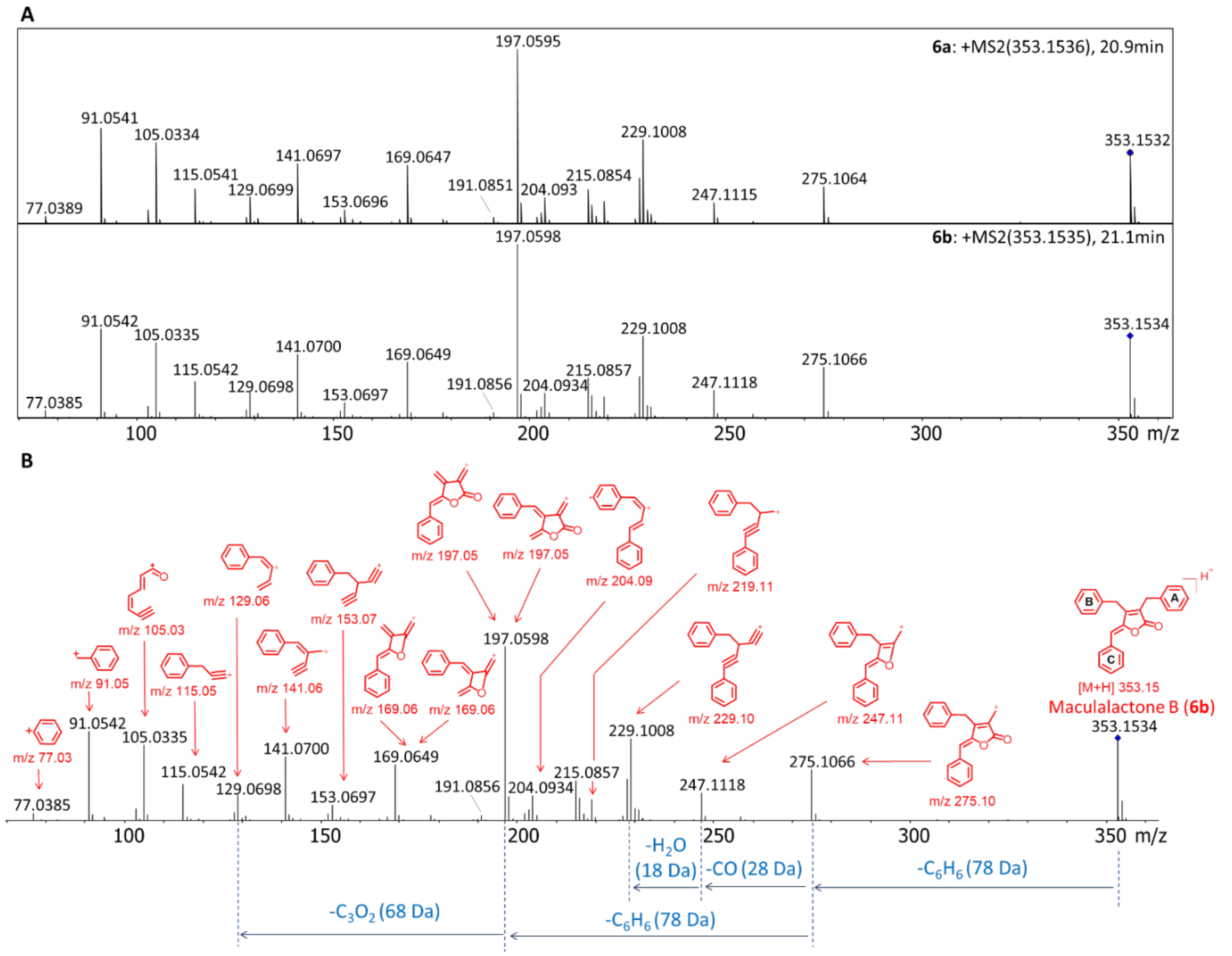


**Figure S9. HRMSMS and fragment ion annotation of NMR elucidated compound 6 with *m/z* 353 $[M+H]^+$: A)** Individual HRMSMS spectra for compound **6b** (maculalactone B) and **6a** (maculalactone C) acquired in ESI positive mode using a data dependent acquisition (DDA) method with a collision energy spread of 20-50 eV. The blue diamond in the spectra indicates the parent ion. **B)** HRMSMS spectrum of **6b** with its respective putative gas phase fragments.

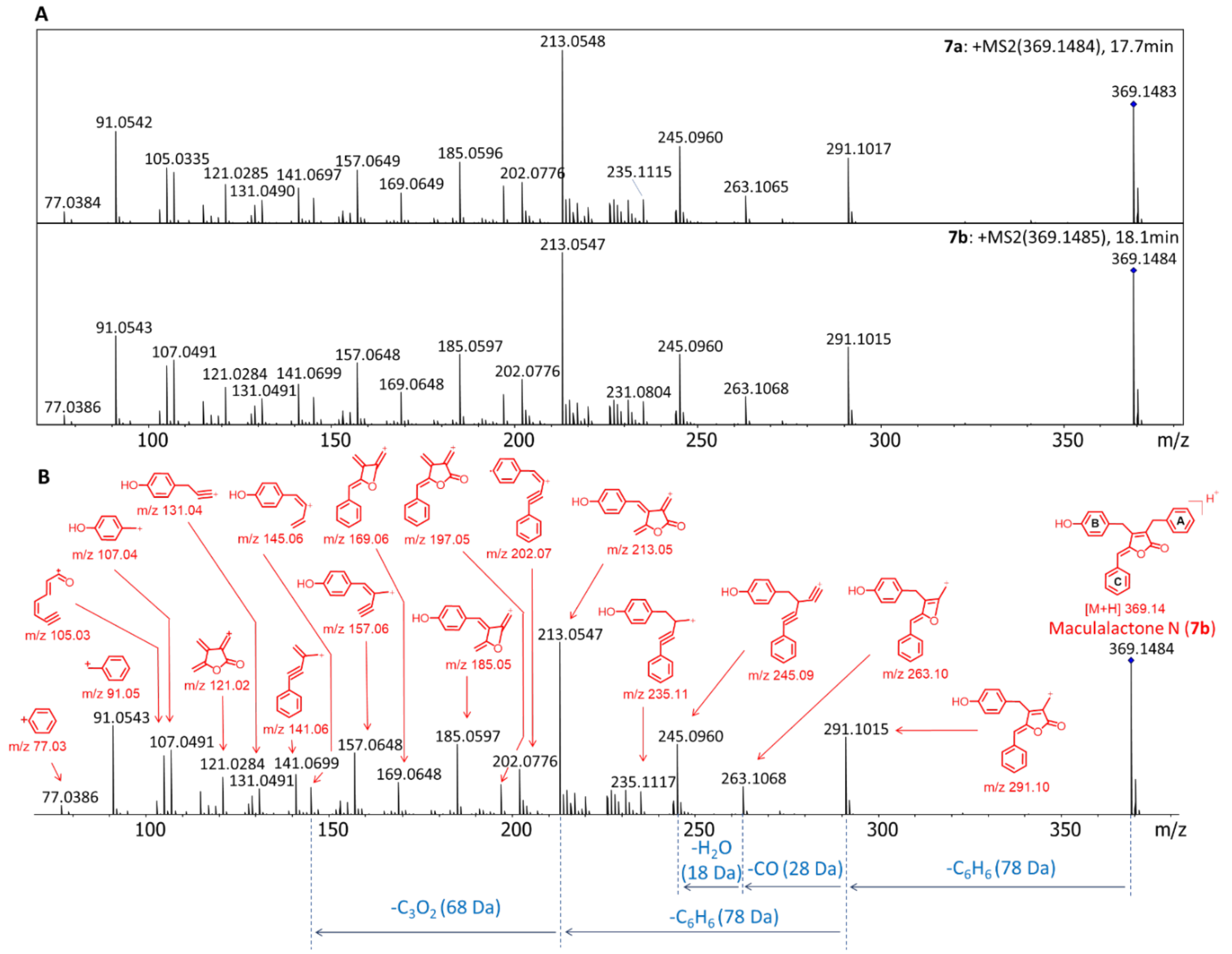


**Figure S10. HRMSMS and fragment ion annotation of NMR elucidated compound 7 with *m/z* 369 [M+H]$^+$: A)** Individual HRMSMS spectra for compound **7b** (maculalactone N) and **7a** acquired in ESI positive mode using a DDA method with a collision energy spread of 20-50 eV. The blue diamond in the spectra indicates the parent ion. **B)** HRMSMS spectrum of **7b** with its respective putative gas phase fragments.

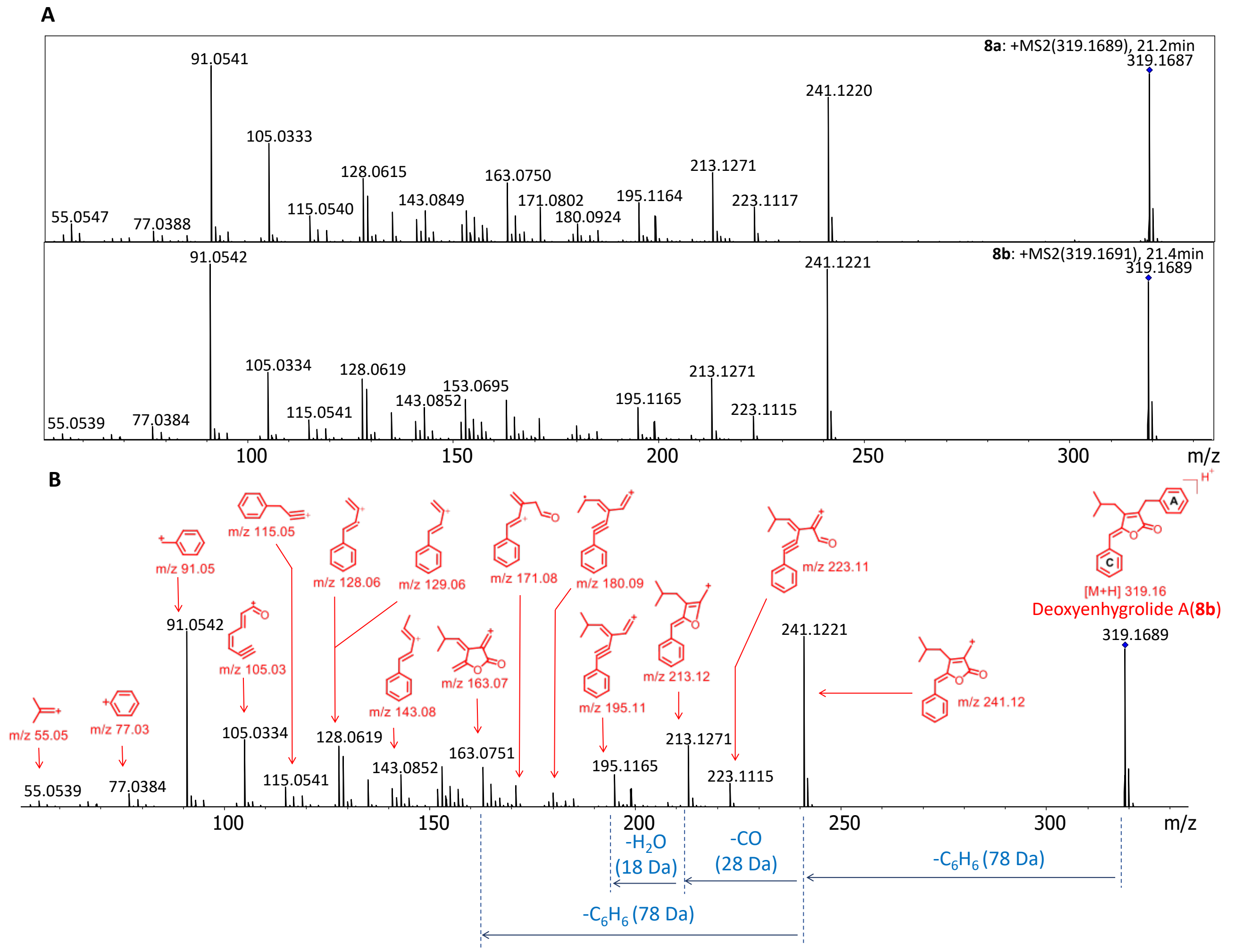


**Figure S11. HRMS/MS and fragment ion annotation of NMR elucidated compound 8 with *m/z* 319 [M+H]$^+$: A)** Individual HRMSMS spectra for compounds **8a** and **8b** acquired in ESI positive mode using a DDA method with a collision energy spread of 20-50 eV. The blue diamond in the spectra indicates the parent ion. **B)** HRMSMS spectrum of **8b** with its respective putative gas phase fragments.

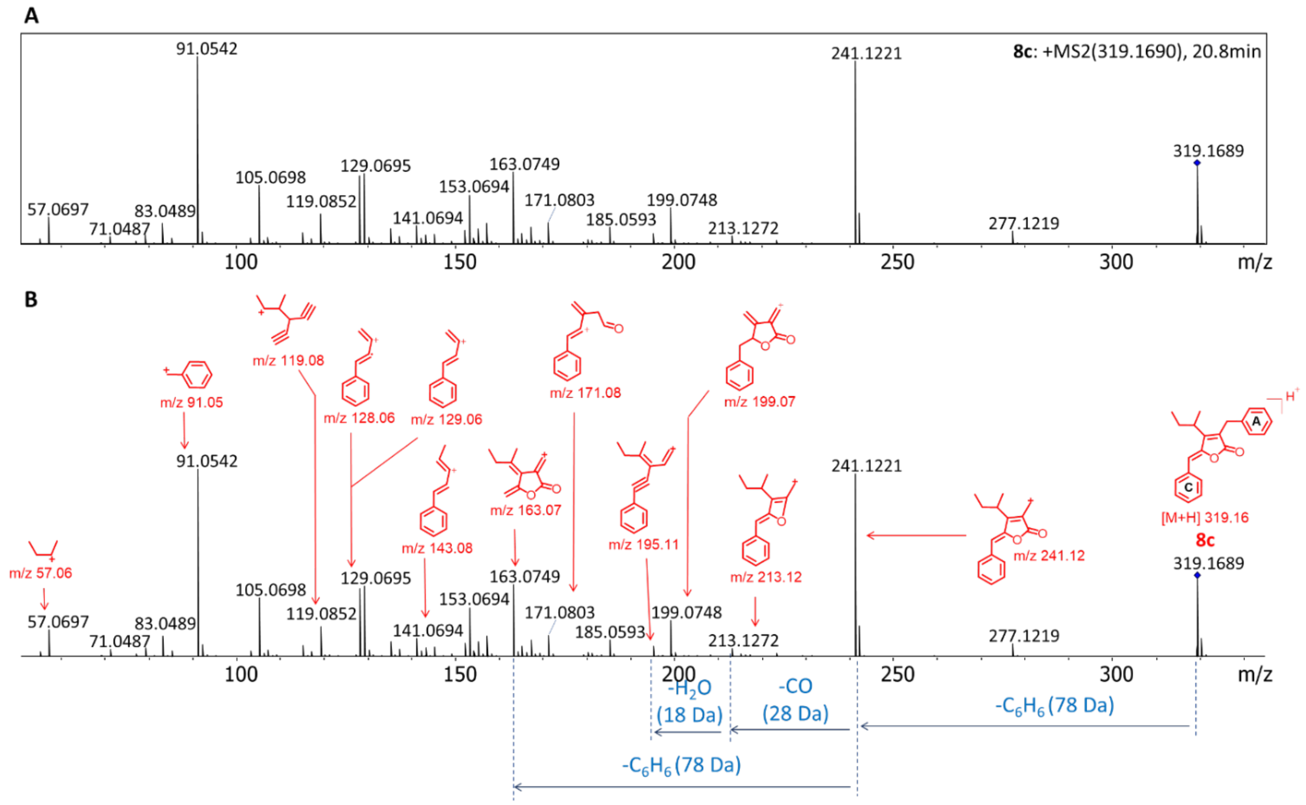


**Figure S12. HRMS/MS and fragment ion annotation of compound 8c with *m/z* 319 $[M+H]^+$: A)** Individual HRMSMS spectra for compound **8c** acquired in ESI positive mode using a DDA method with a collision energy spread of 20-50 eV. The blue diamond in the spectra indicates the parent ion. **B)** HRMSMS spectrum of **8c** with its respective putative gas phase fragments.

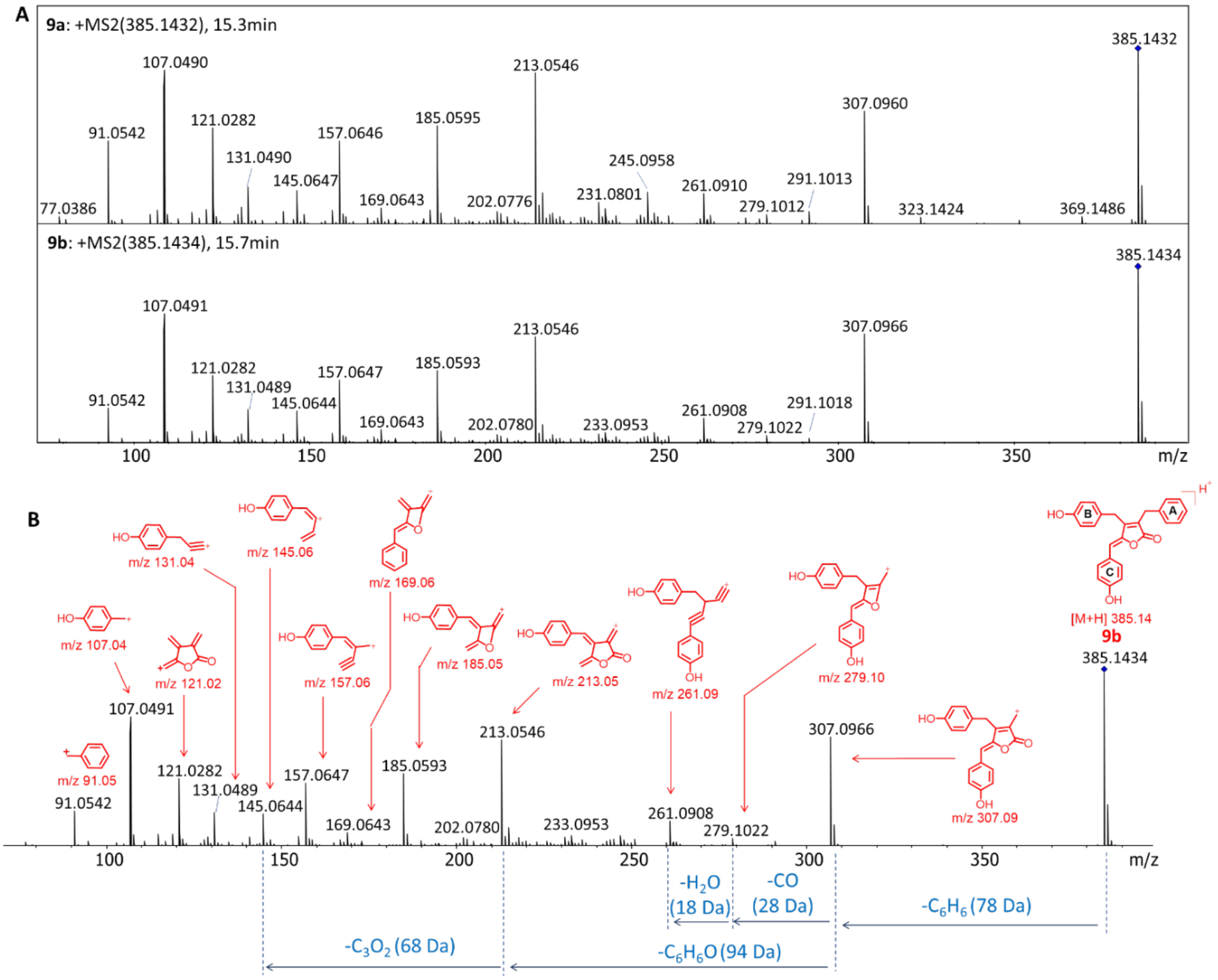


**Figure S13. HRMS/MS and fragment ion annotation of *m/z* 385 $[M+H]^+$: A)** Individual HRMSMS spectra for the phenol mono-hydroxylated compounds **9a** and **9b** acquired in ESI positive mode using a DDA method with a collision energy spread of 20-50 eV. The blue diamond in the spectra indicates the parent ion. **B)** HRMSMS spectrum of compound **9b** with its respective putative gas phase fragments.

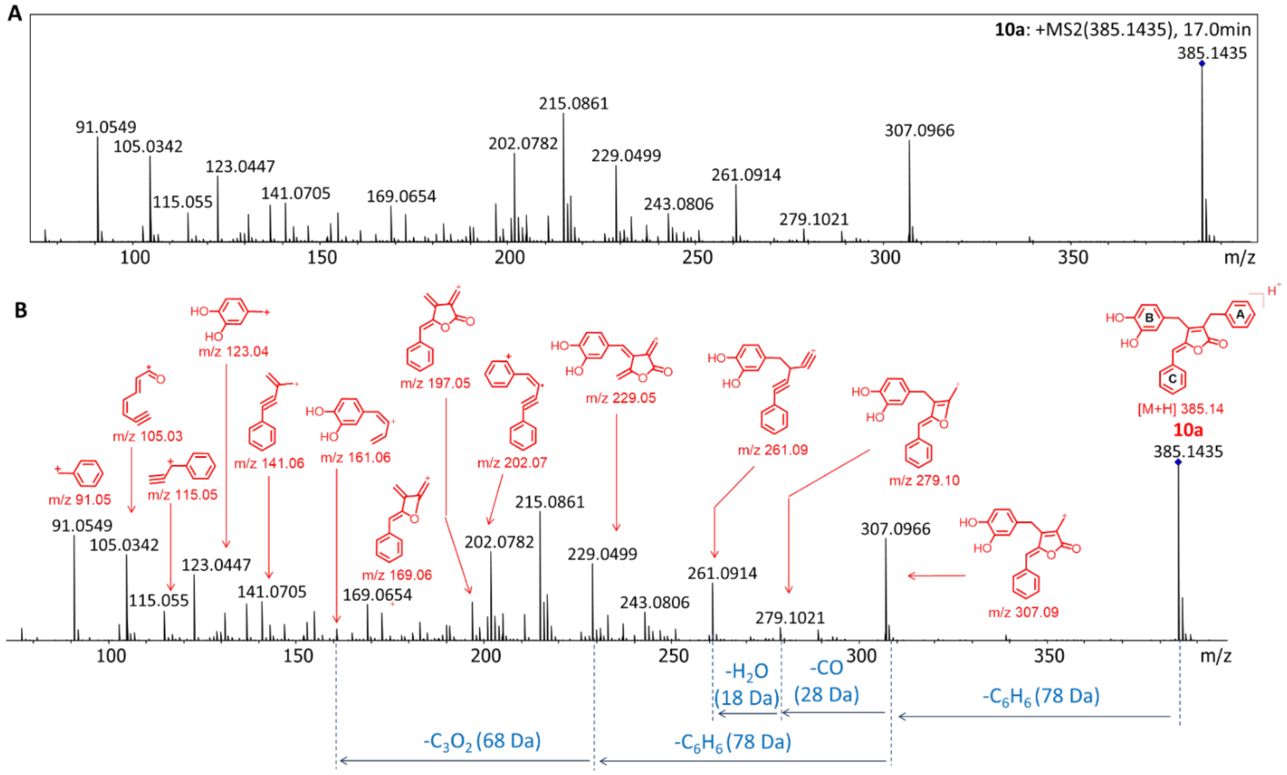


**Figure S14. HRMSMS and fragment ion annotation of *m/z* 385 [M+H]$^+$: A)** Individual HRMSMS spectra for phenol dihydroxylated compound **10a** acquired in ESI positive mode using a DDA method with a collision energy spread of 20-50 eV. The blue diamond in the spectra indicates the parent ion. **B)** HRMSMS spectrum of **10a** with its respective putative gas phase fragments.

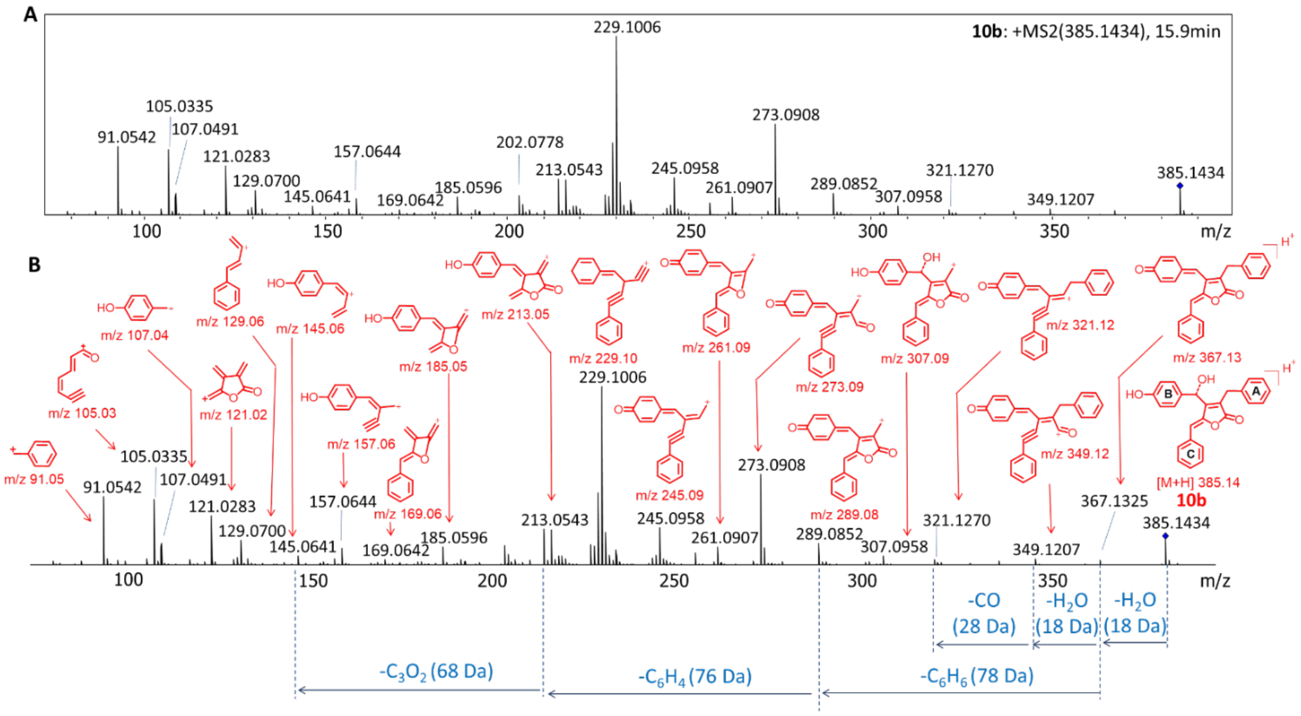


**Figure S15. HRMSMS and fragment ion annotation of *m/z* 385 [M+H]$^+$: A)** Individual HRMSMS spectra for compound **10b** acquired in ESI positive mode using a DDA method with a collision energy spread of 20-50 eV. The blue diamond in the spectra indicates the parent ion. **B)** HRMSMS spectrum of **10b** with its respective putative gas phase fragments.

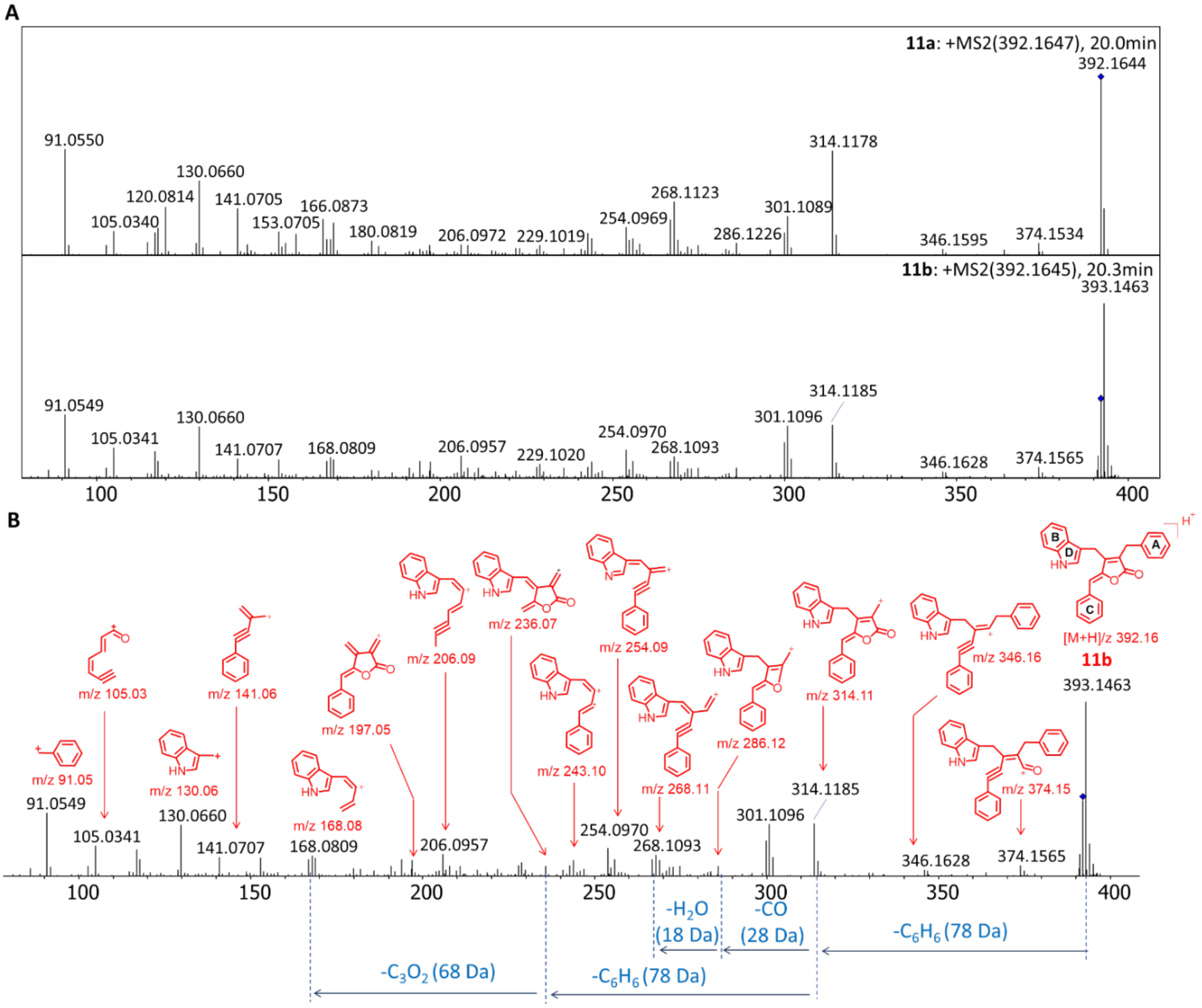


**Figure S16. HRMSMS and fragment ion annotation of *m/z* 392 [M+H]$^+$: A)** Individual HRMS/MS spectra for compounds **11a** and **11b** acquired in ESI positive mode using a DDA with a collision energy spread of 20-50 eV. The blue diamond in the spectra indicates the parent ion. **B)** HRMSMS spectrum of **11b** with its respective putative gas phase fragments.

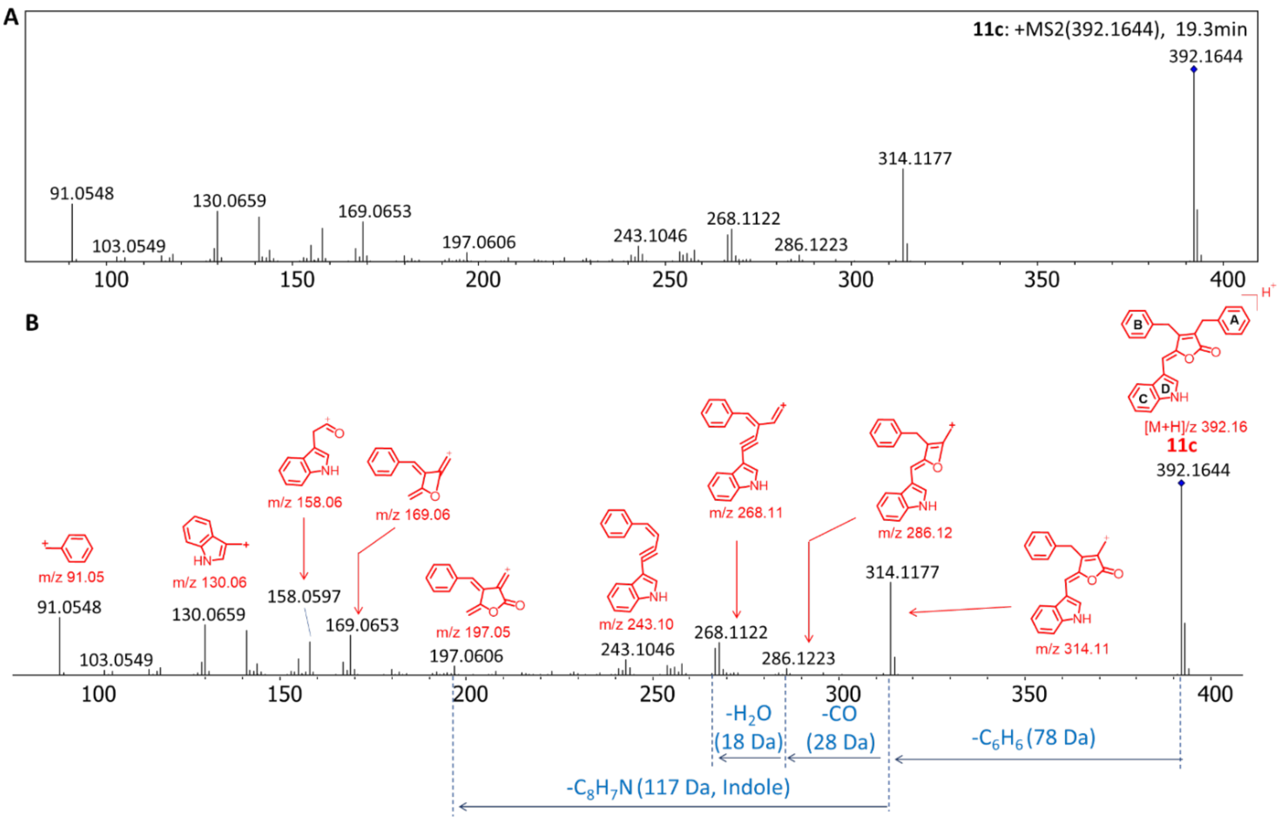


**Figure S17. HRMSMS and fragment ion annotation of *m/z* 392 [M+H]$^+$: A)** Individual HRMS/MS spectrum for compounds **11c** acquired in ESI positive mode using a DDA with a collision energy spread of 20-50 eV. The blue diamond in the spectra indicates the parent ion. **B)** HRMSMS spectrum of **11c** with its respective putative gas phase fragments.

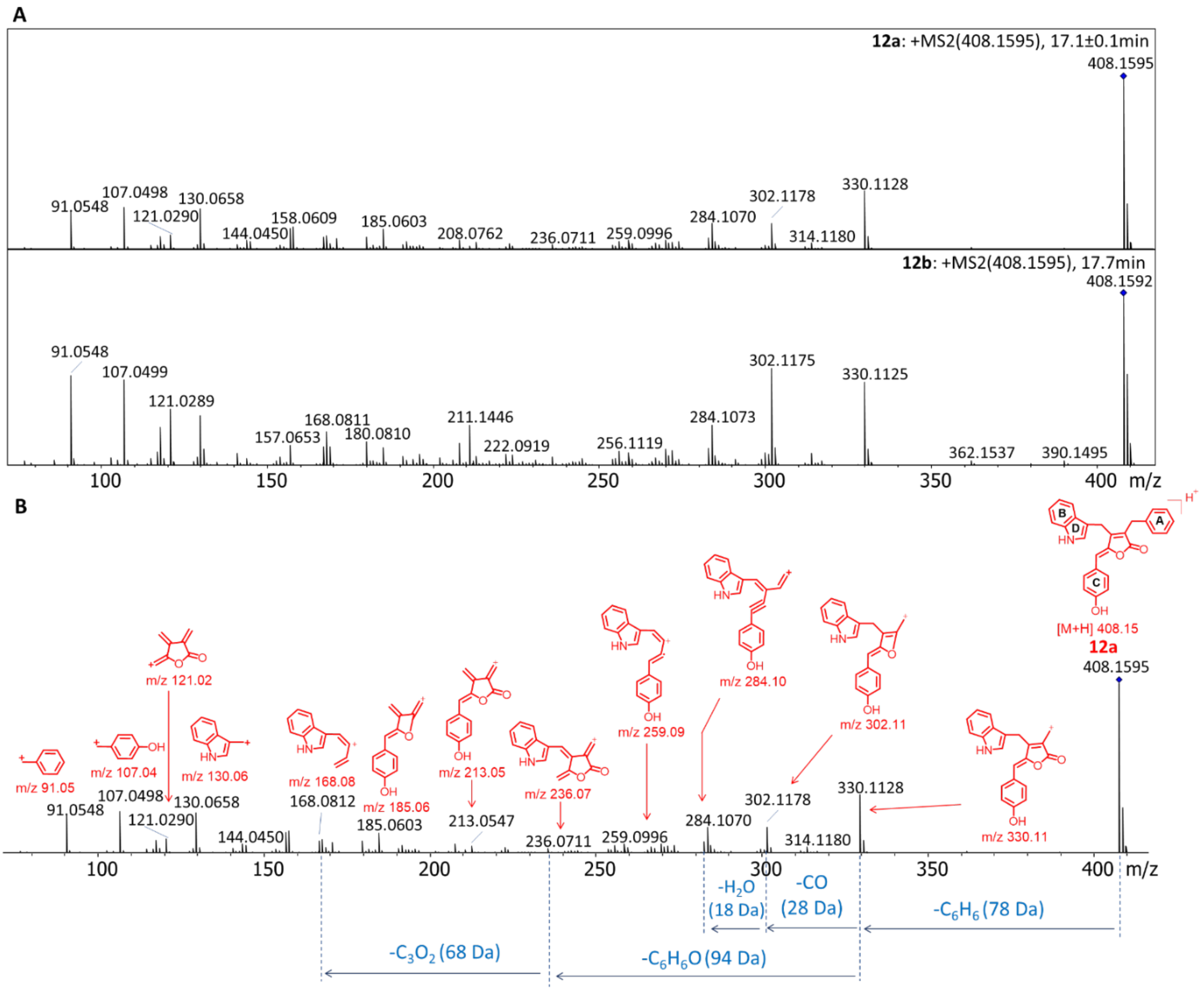


**Figure S18. HRMSMS and fragment ion annotation of *m/z* 408 [M+H]$^+$: A)** Individual HRMSMS spectra for compounds **12a** and **12b** acquired in ESI positive mode using a DDA method with a collision energy spread of 20-50 eV. The blue diamond in the spectra indicates the parent ion. **B)** HRMSMS spectrum of **12b** with its respective putative gas phase fragments.

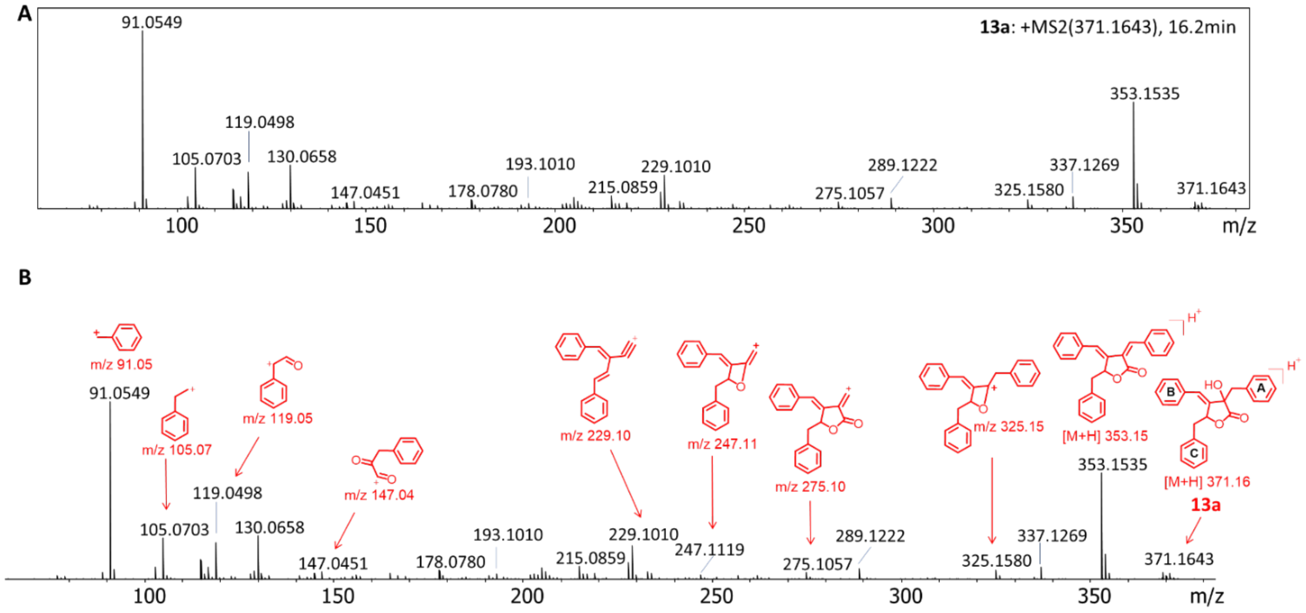


**Figure S19. HRMSMS and fragment ion annotation of *m/z* 371 $[M+H]^+$: A)** Individual HRMSMS spectrum for compound **13a** (maculalactone L) acquired in ESI positive mode using a DDA method with a collision energy spread of 20-50 eV. The blue diamond in the spectra indicates the parent ion. **B)** HRMSMS spectrum of **13a** with its respective putative gas phase fragments.

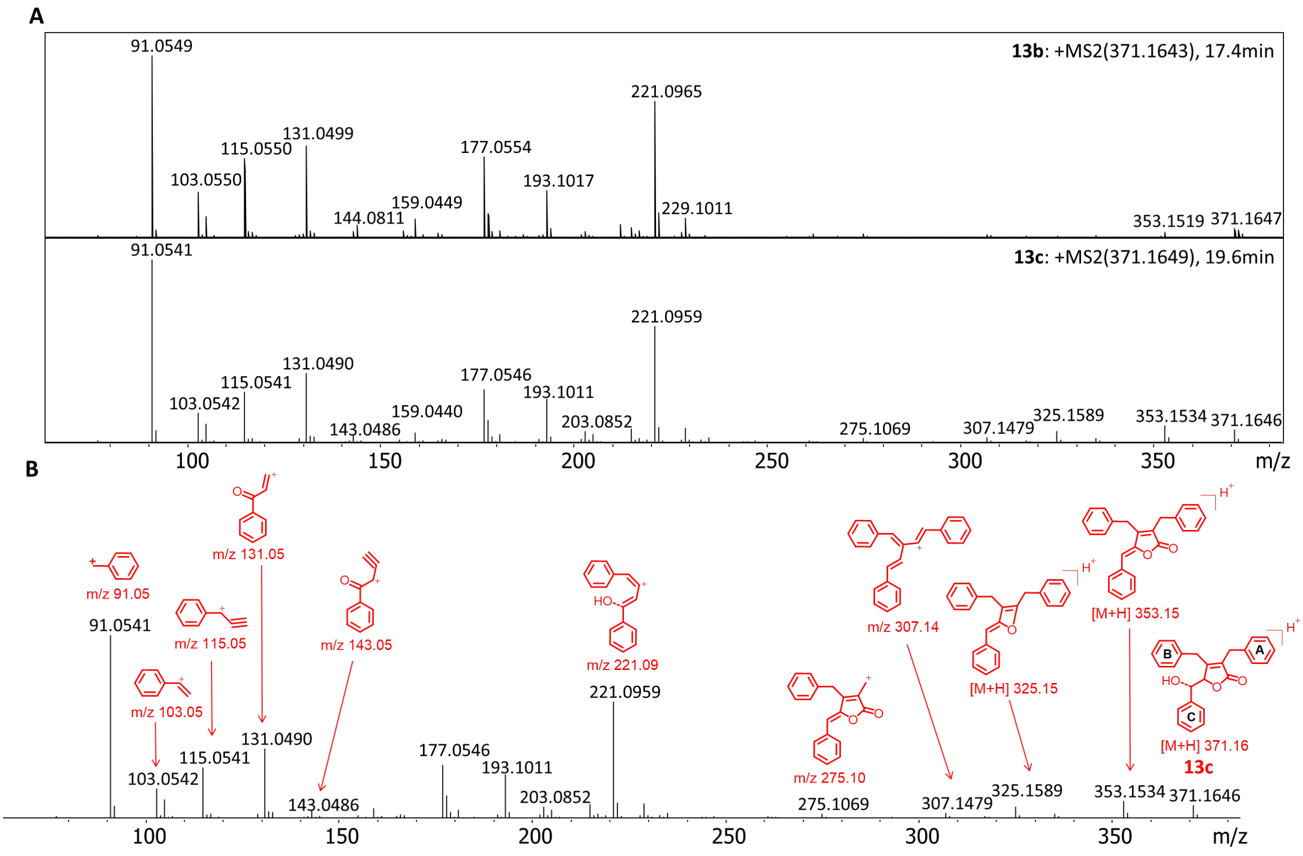


**Figure S20. HRMSMS and fragment ion annotation of *m/z* 371 $[M+H]^+$: A)** Individual HRMSMS spectra for compounds **13b** and **13c** acquired in ESI positive mode using a DDA method with a collision energy spread of 20-50 eV. The blue diamond in the spectra indicates the parent ion. **B)** HRMSMS spectrum of **13c** with its respective putative gas phase fragments.

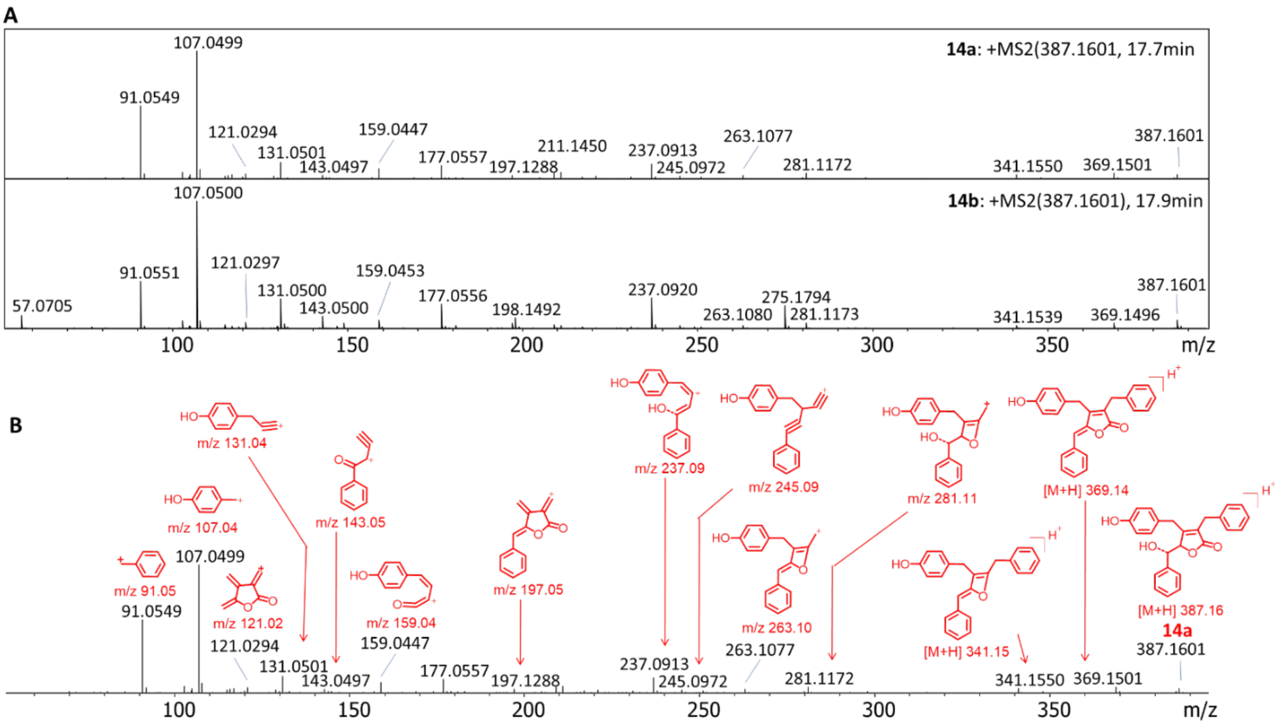


**Figure S21. HRMSMS and fragment ion annotation of *m/z* 387 [M+H]$^+$: A)** Individual HRMS/MS spectra for compounds **14a** and **14b** acquired in ESI positive mode using a DDA method with a collision energy spread of 20-50 eV. The blue diamond in the spectra indicates the parent ion. **B)** HRMSMS spectrum of **14a** with its respective putative gas phase fragments.

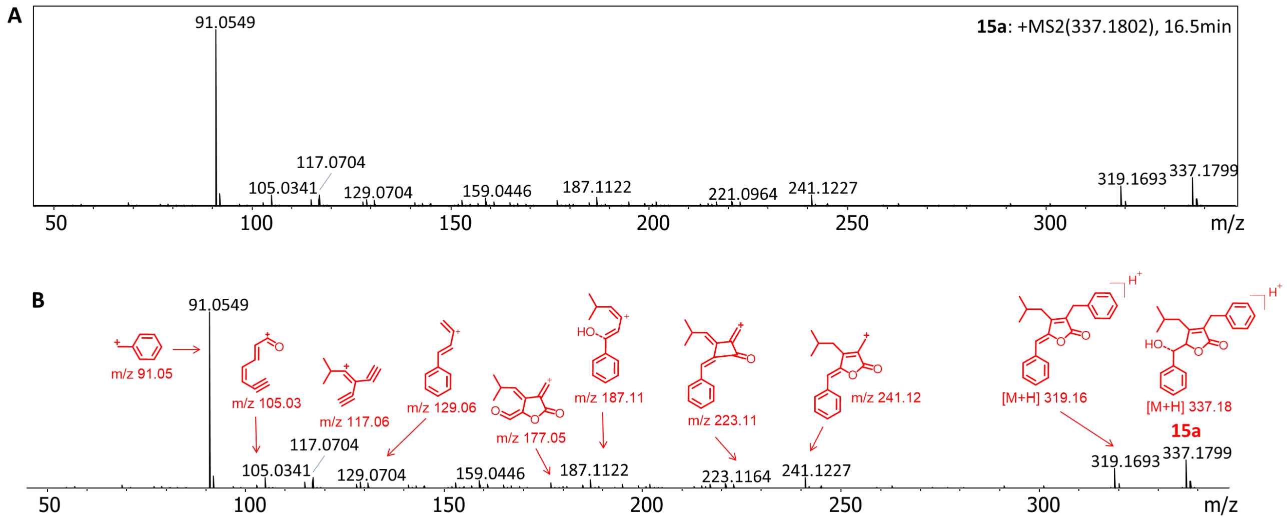


**Figure S22. HRMSMS and fragment ion annotation of *m/z* 337 [M+H]$^{+}$: A)** Individual HRMS/MS spectrum for compound **15a** acquired in ESI positive mode using a DDA method with a collision energy spread of 20-50 eV. The blue diamond in the spectra indicates the parent ion. **B)** HRMSMS spectrum of **15a** with its respective putative gas phase fragments

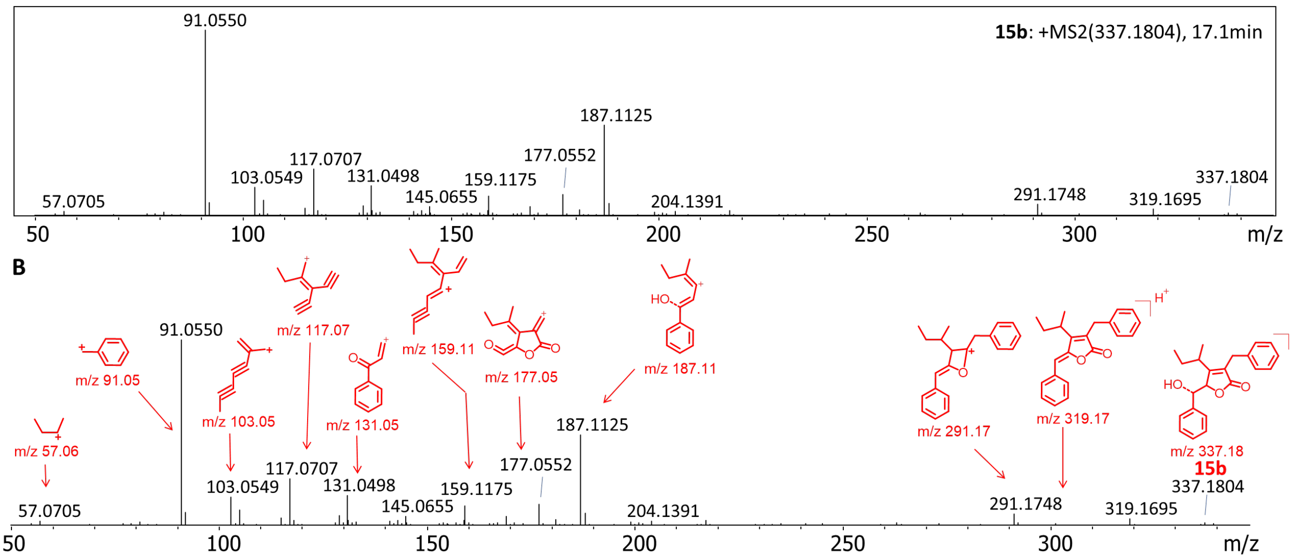


**Figure S23. HRMSMS and fragment ion annotation of *m/z* 337 [M+H]$^{+}$: A)** Individual HRMS/MS spectrum for compound **15b** acquired in ESI positive mode using a DDA method with a collision energy spread of 20-50 eV. The blue diamond in the spectra indicates the parent ion. **B)** HRMSMS spectrum of **15b** with its respective putative gas phase fragments

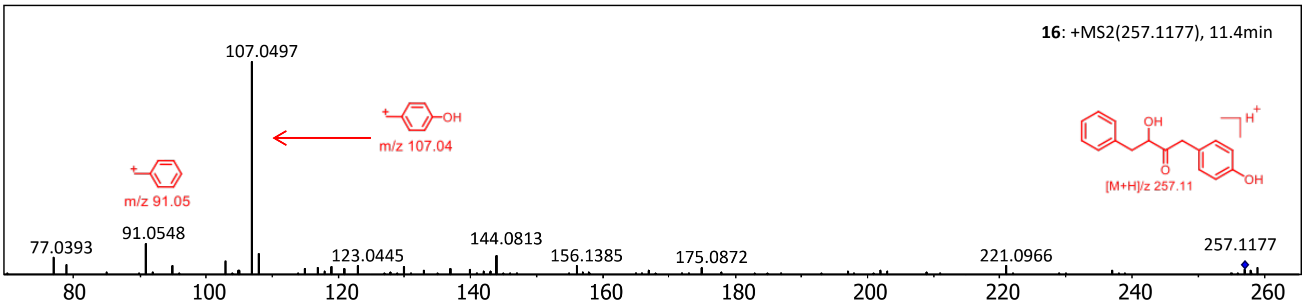


**Figure S24. HRMSMS and fragment ion annotation of the known acyloin kurasoin A/B (*m/z* 337 $[M+H]^+$):** Kurasoin A/B (**16**) is an acyloin compound and the likely product of the TPP-dependant enzyme MacE.

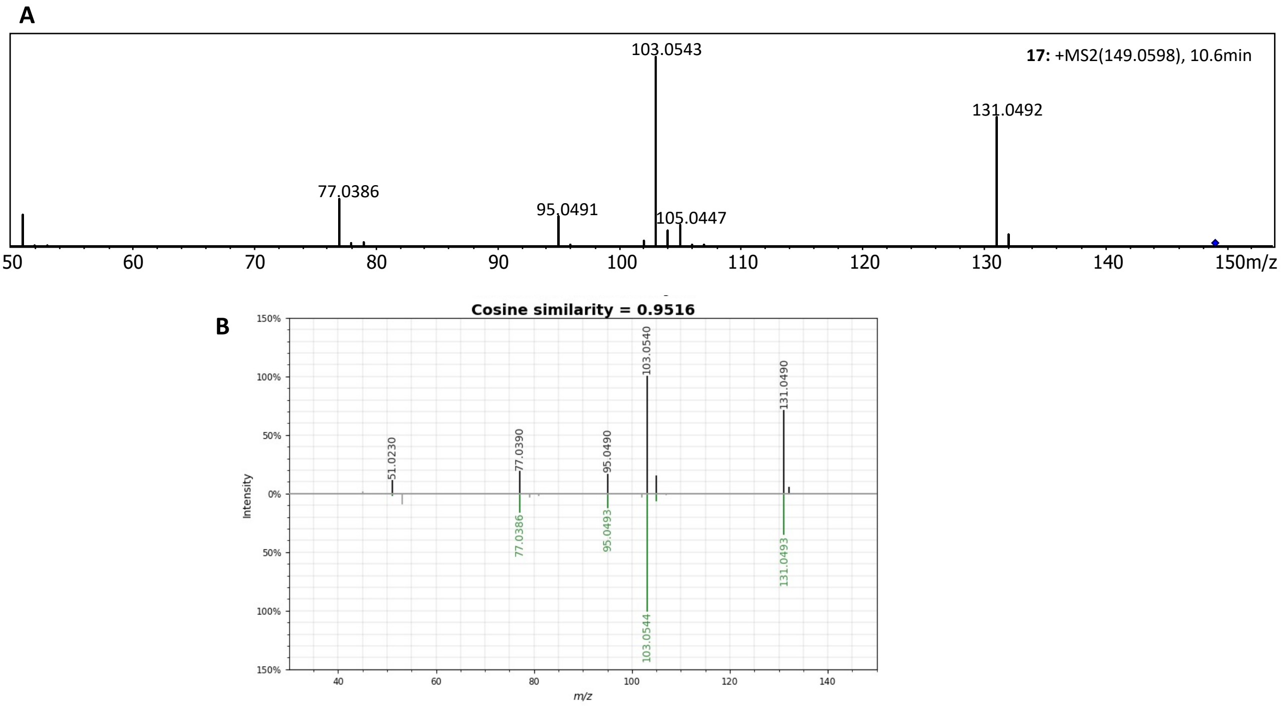


**Figure S25. HRMSMS data of cinnamic acid (17): A)** Individual HRMSMS spectrum of compound **17** acquired in ESI positive mode using a DDA method with a collision energy spread of 20-50 eV. The blue diamond in the spectra indicates the parent ion. **B)** Mirror image of the MS/MS spectral match (cosine > 0.9) of cinnamic acid between experimental (query) tandem MS spectrum and GNPS library spectrum. The image was generated using USI Resolution tool within GNPS environment.

### NMR spectra

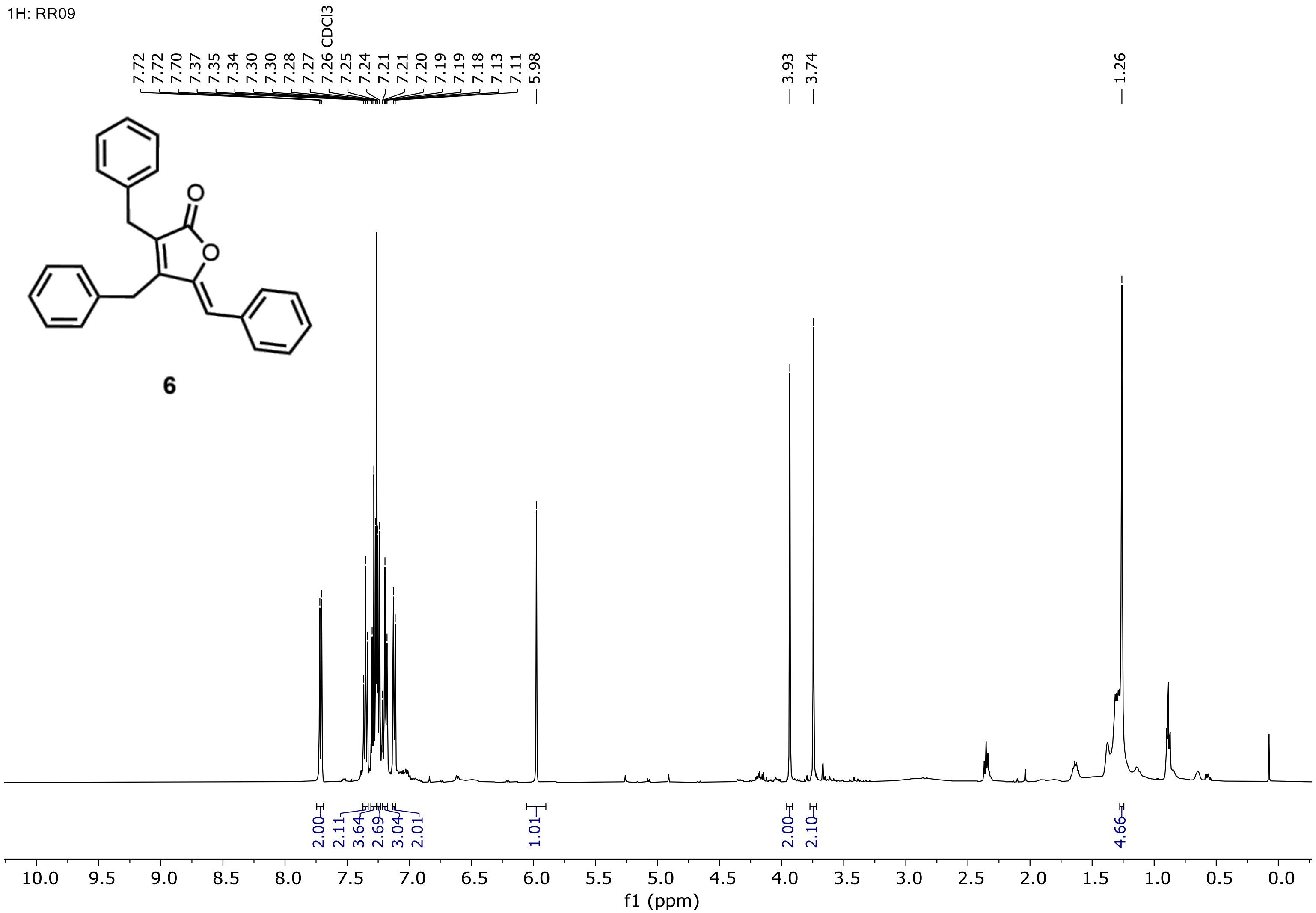


**Figure S26.** $^{1}$H NMR of maculalactone B (6b), measured in $CDCl_3$ at 500 MHz.

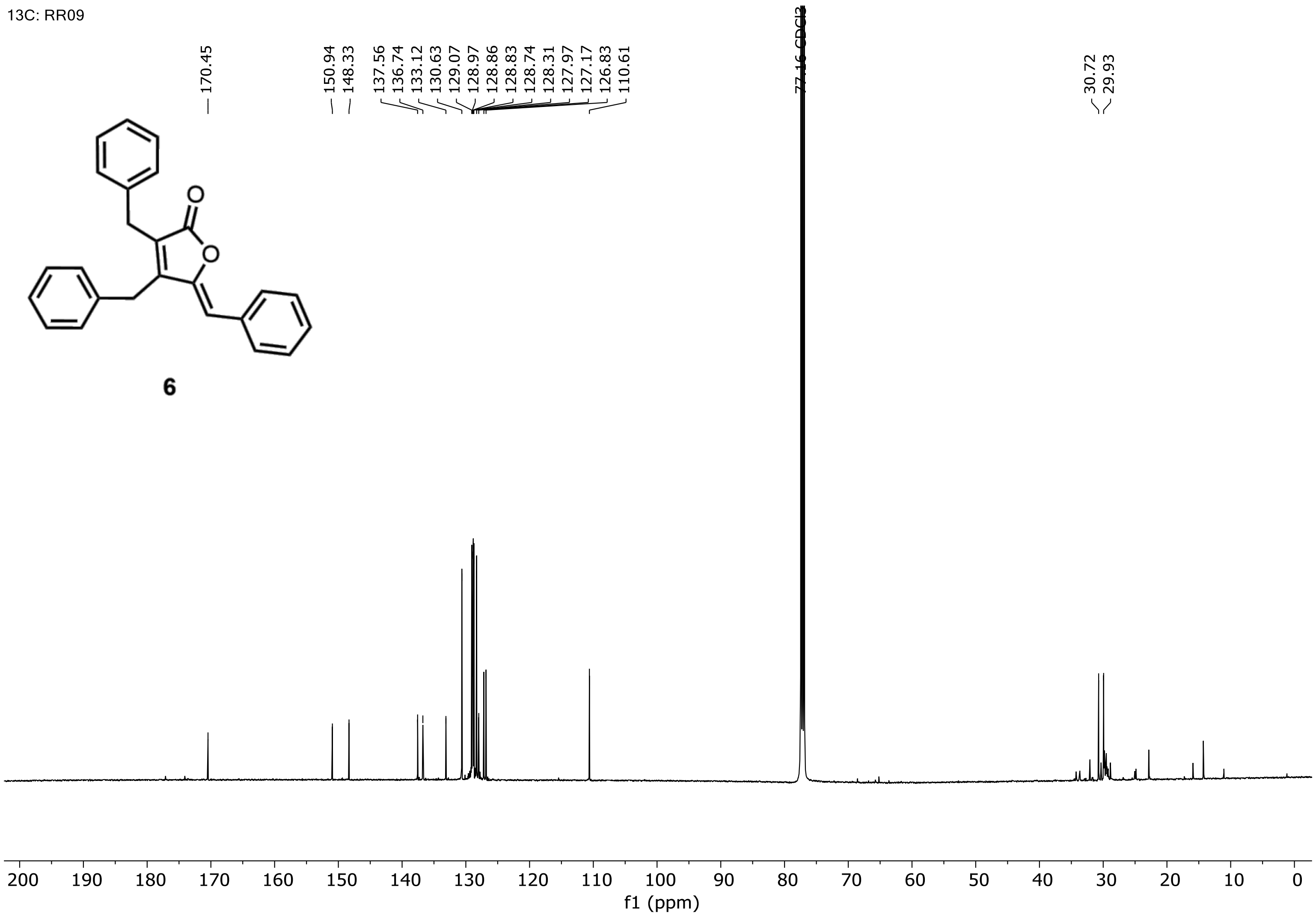


**Figure S27.** **[13]C NMR of maculalactone B (6b), measured in $CDCl_3$ at 126 MHz.**

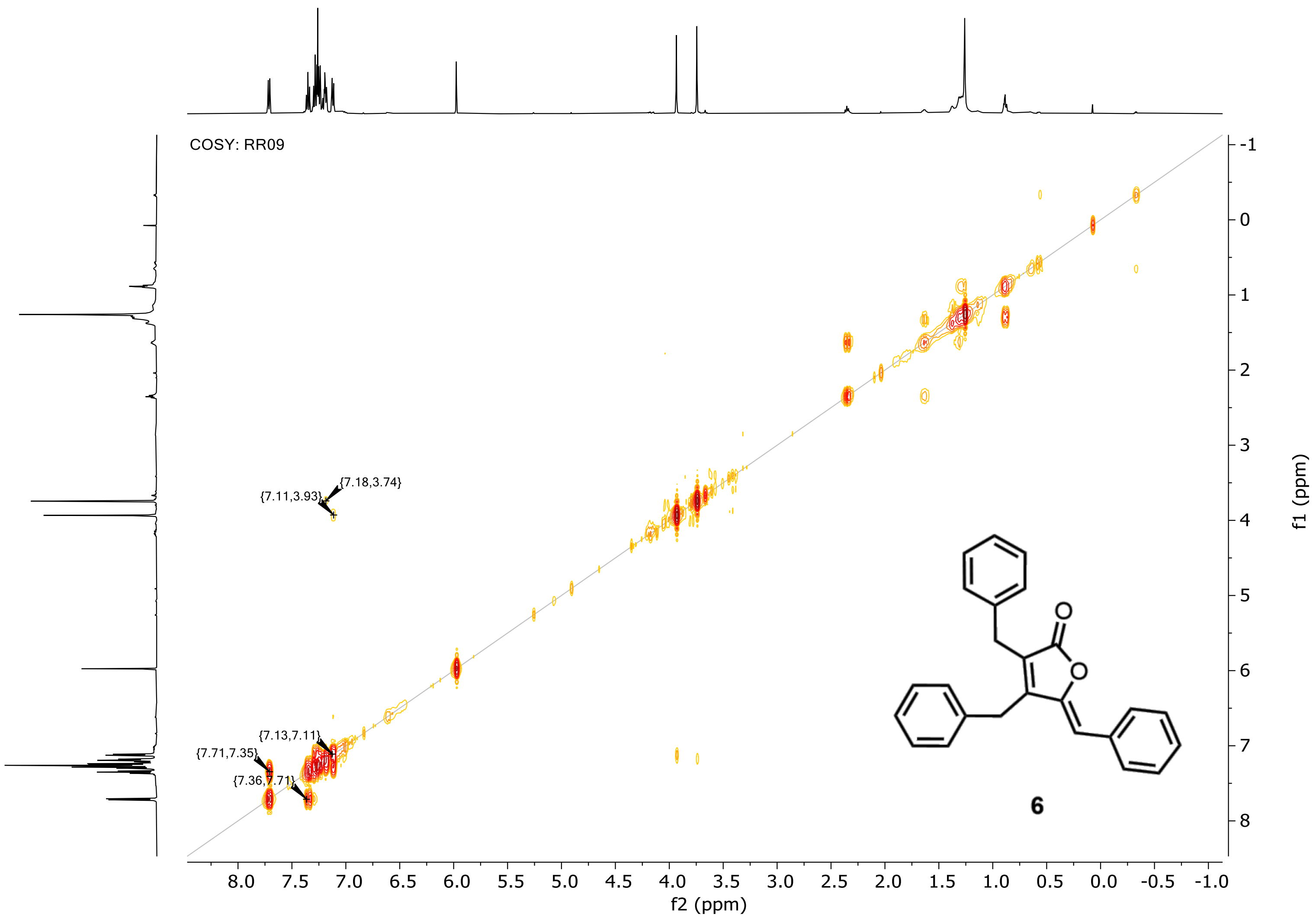


**Figure S28. COSY of maculalactone B (6b), measured in $CDCl_3$.**

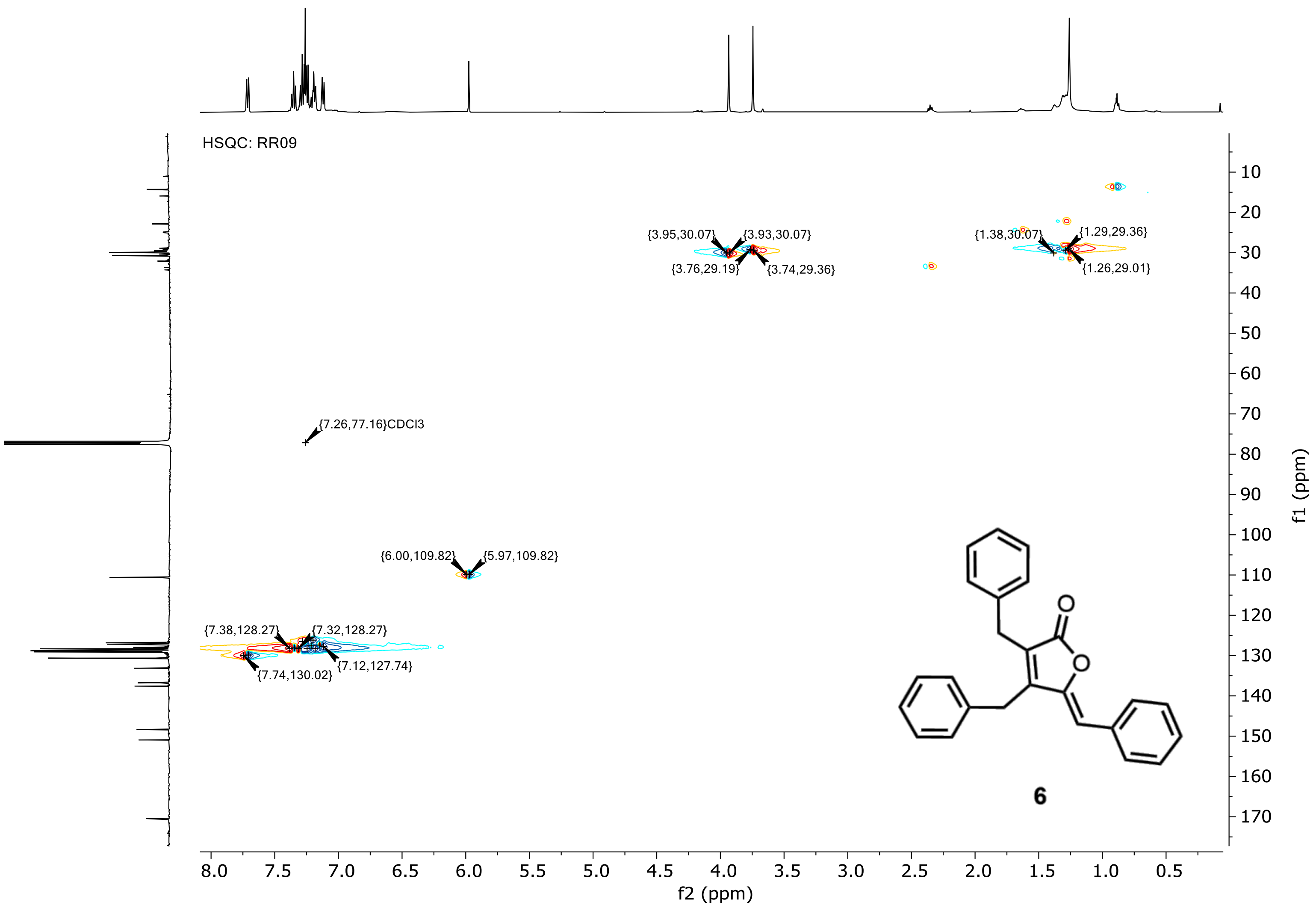


**Figure S29. HSQC of maculalactone B (6b), measured in $CDCl_3$.**

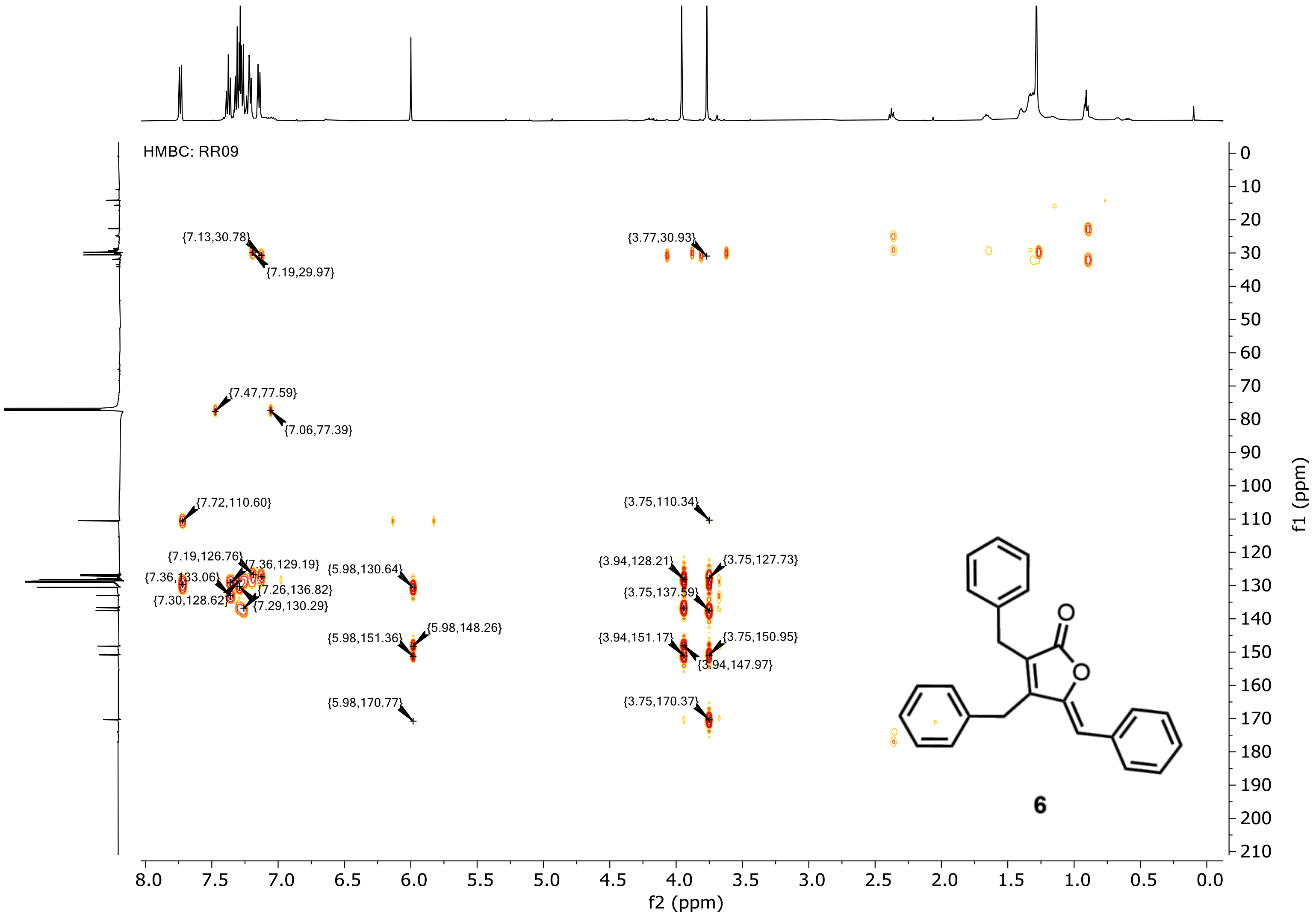


**Figure S30. HMBC of maculalactone B (6b), measured in CDCl$_3$.**

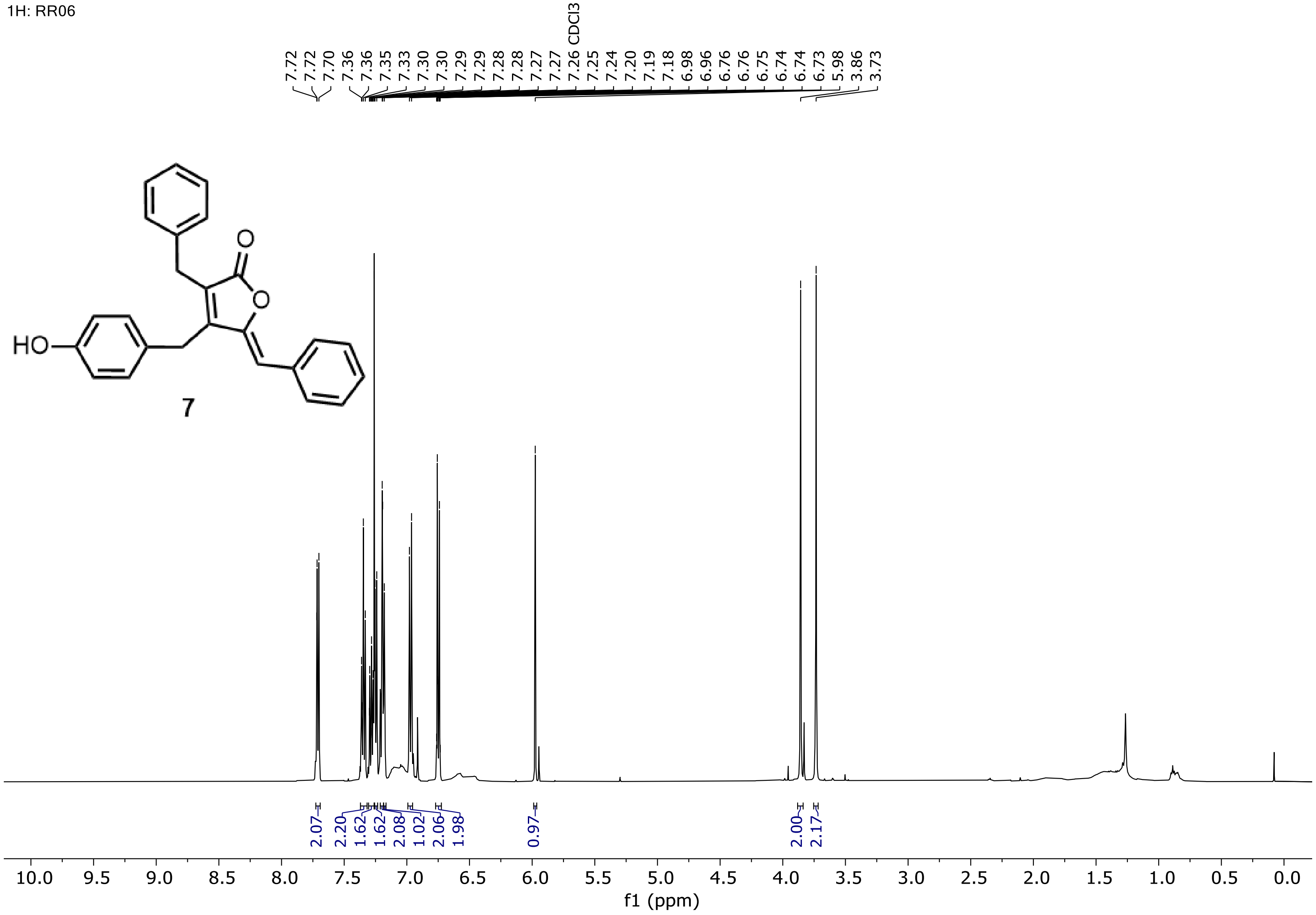


**Figure S31. ¹H NMR of maculalactone N (7b), measured in $CDCl_3$ at 500 MHz.**

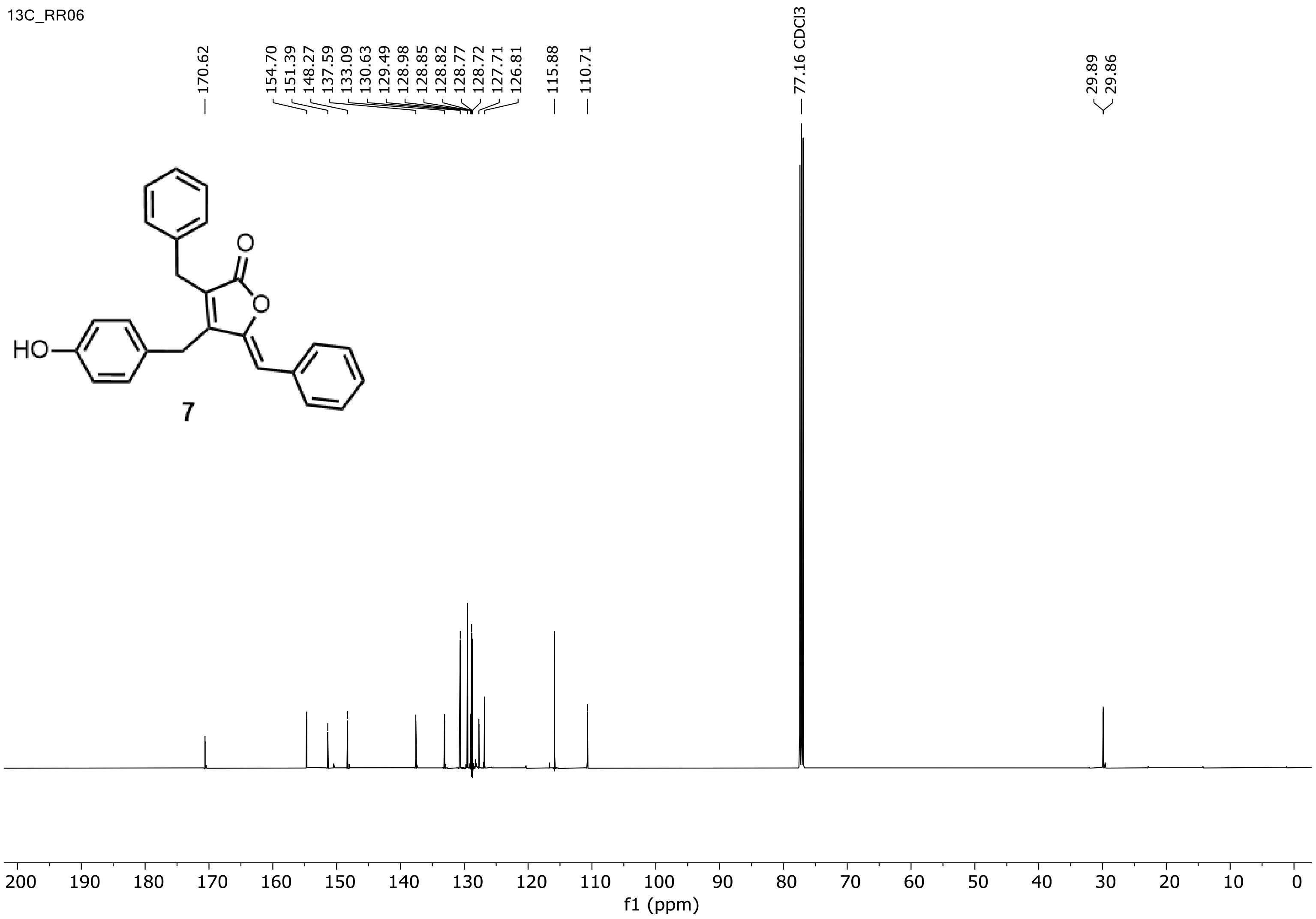


**Figure S32. $^{13}$C NMR of maculalactone N (7b), measured in $CDCl_3$ at 126 MHz.**

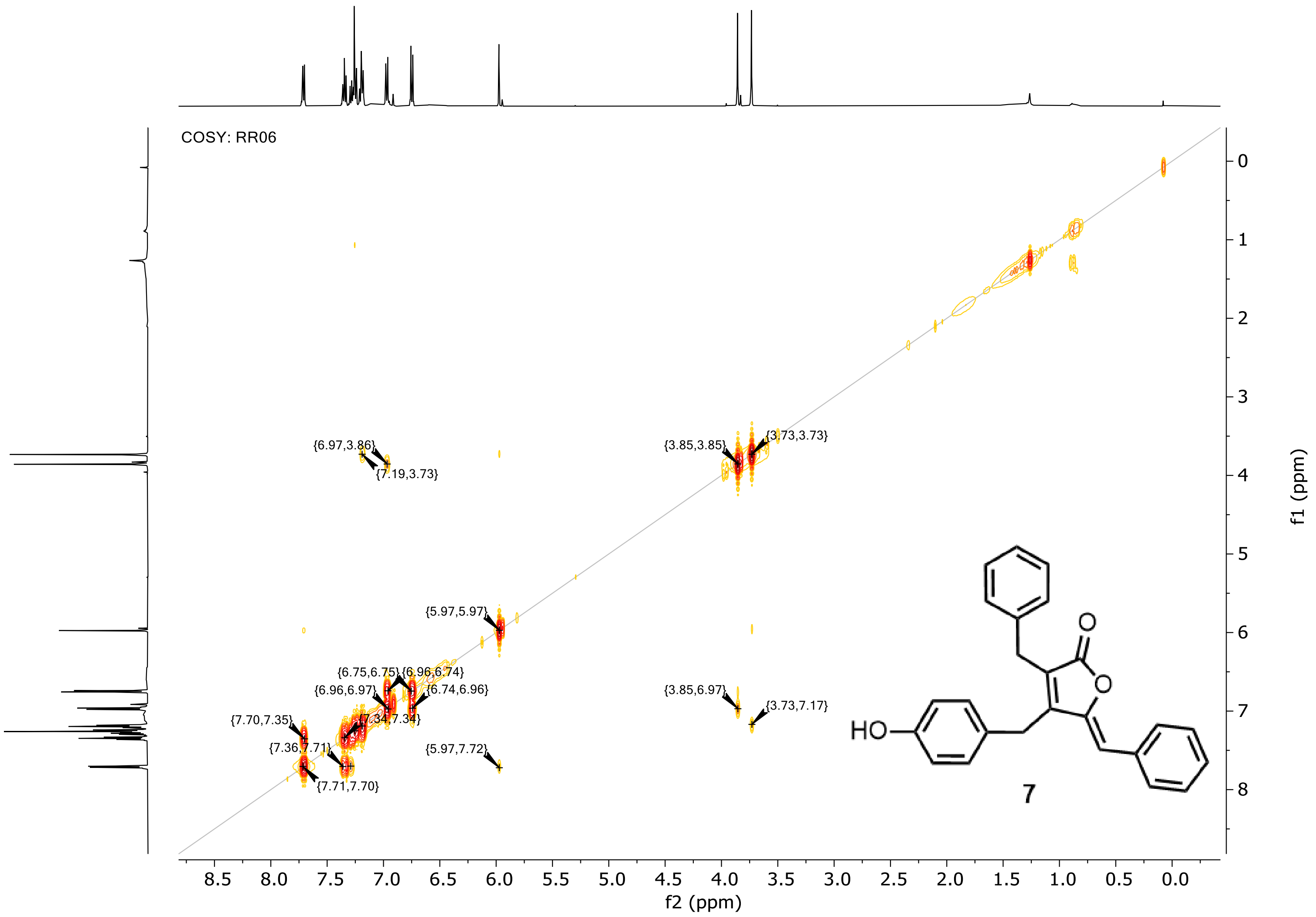


**Figure S33. COSY of maculalactone N (7b), measured in $CDCl_3$.**

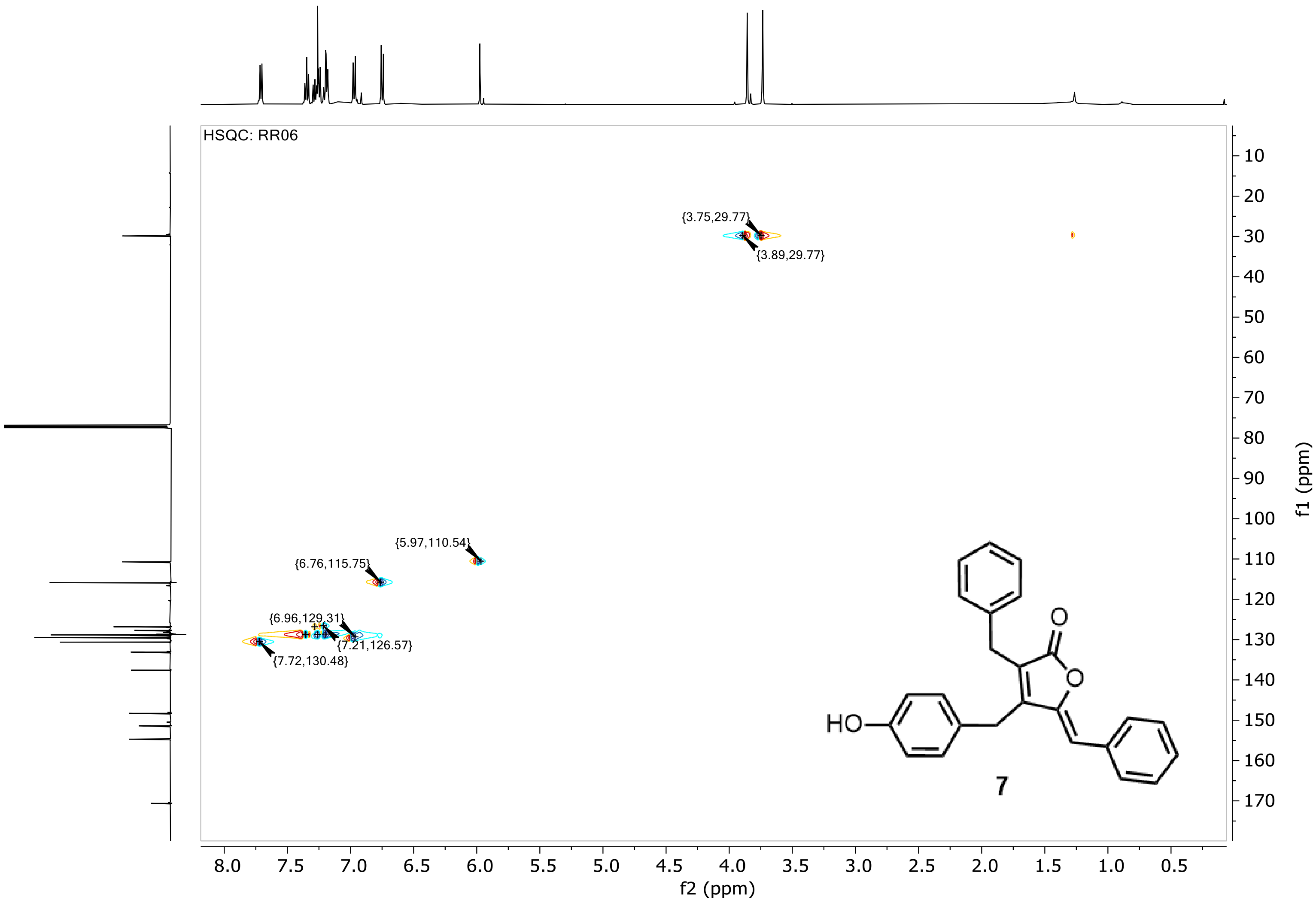


**Figure S34. HSQC of maculalactone N (7b), measured in $CDCl_3$.**

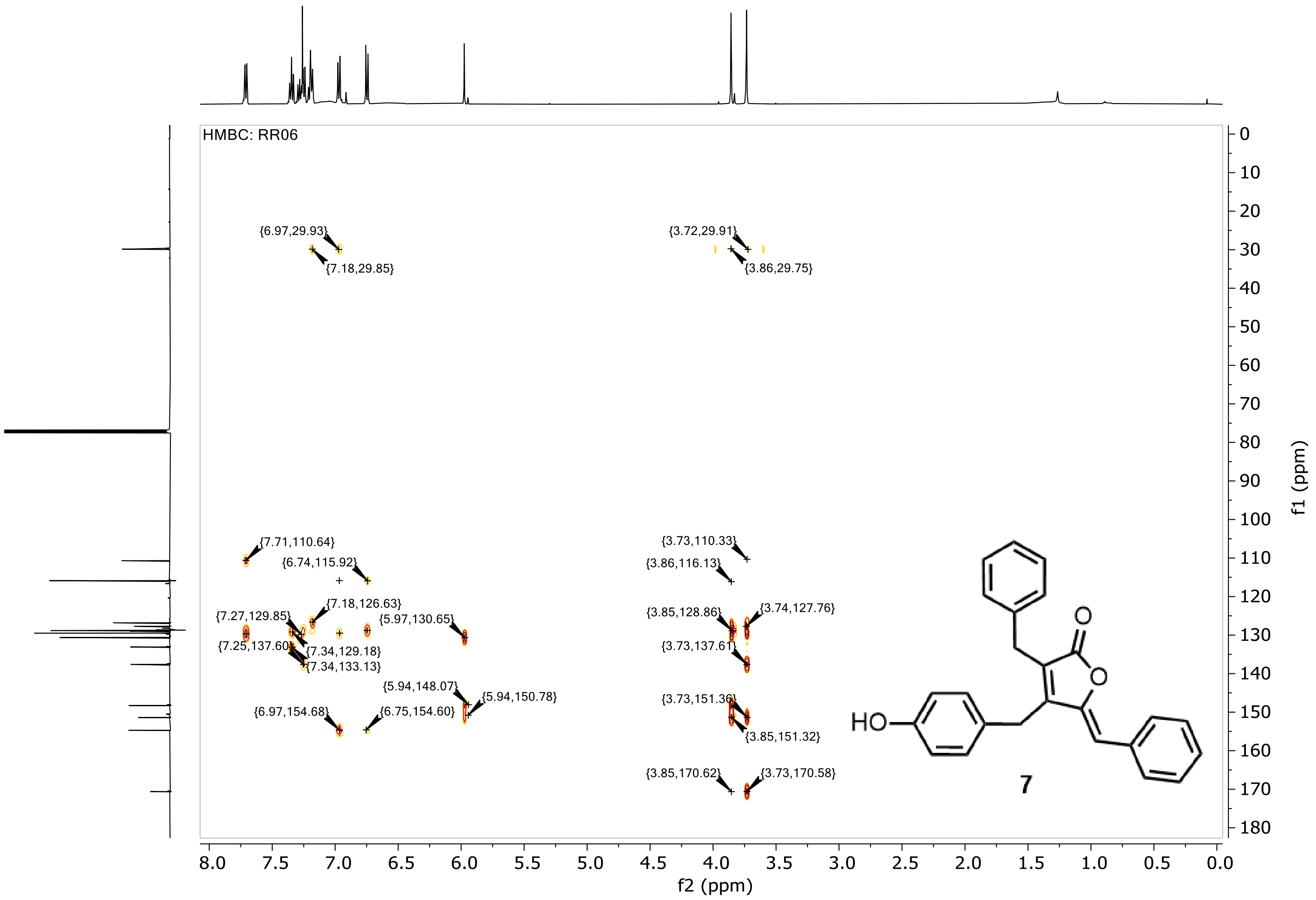


**Figure S35. HMBC of maculalactone N (7b), measured in $CDCl_3$.**

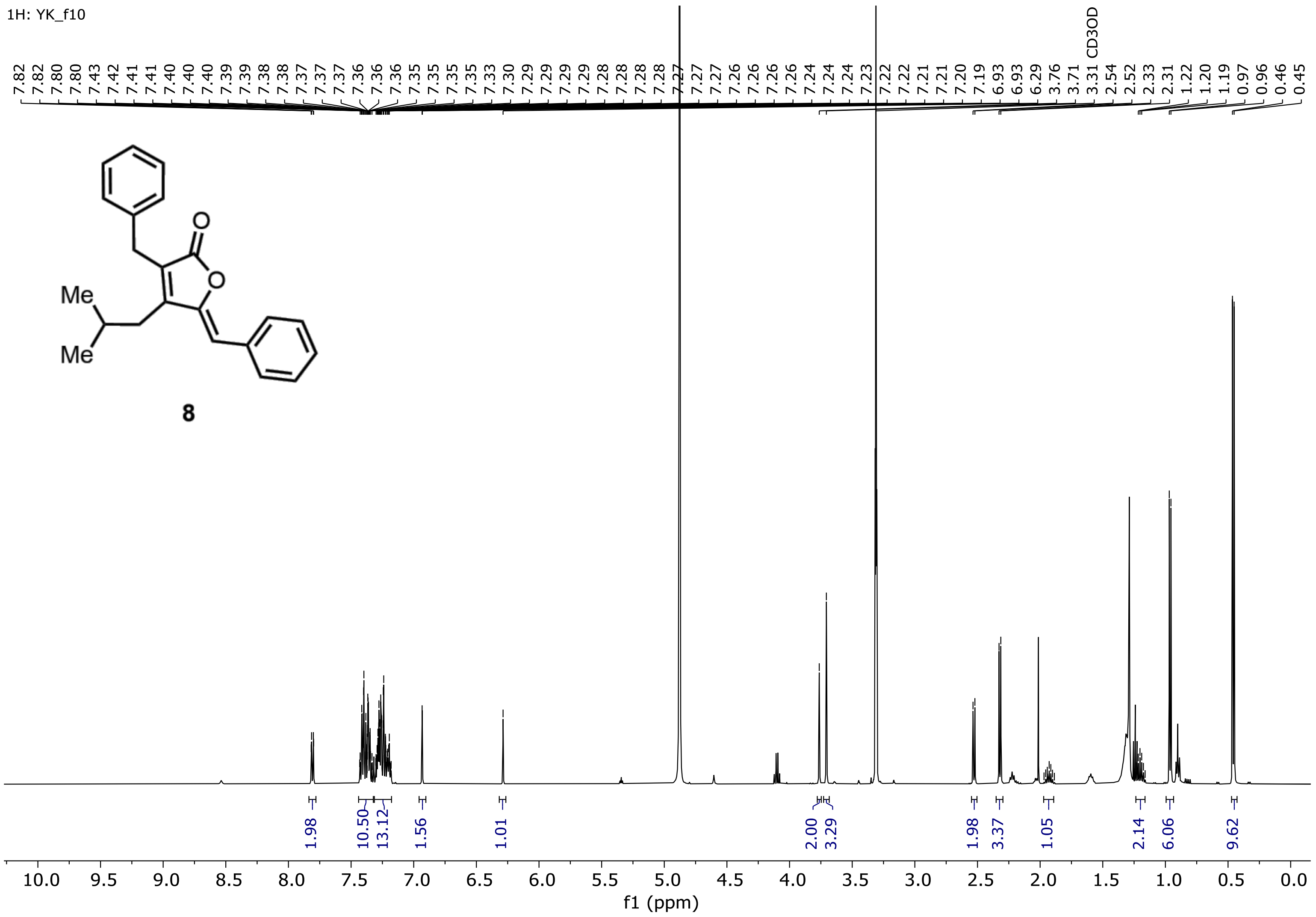


**Figure S36. $^1$H NMR of deoxyenhygrolide A (8b), measured in MeOD-$d_4$ at 500 MHz.**

13C: YK_f10

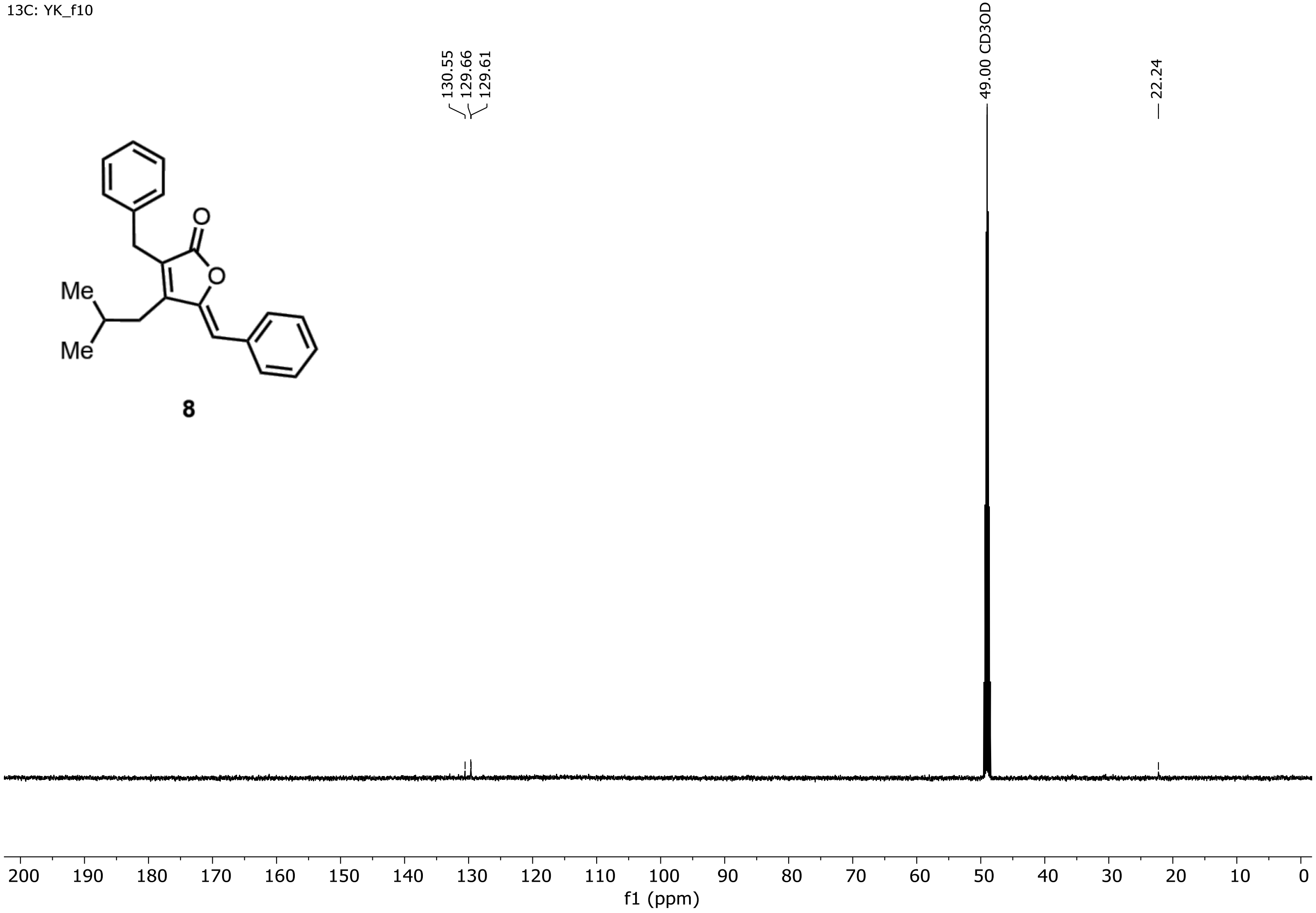


**Figure S37. $^{13}$C NMR of deoxyenhygrolide A (8b), measured in MeOD-$d_4$ at 126 MHz.**

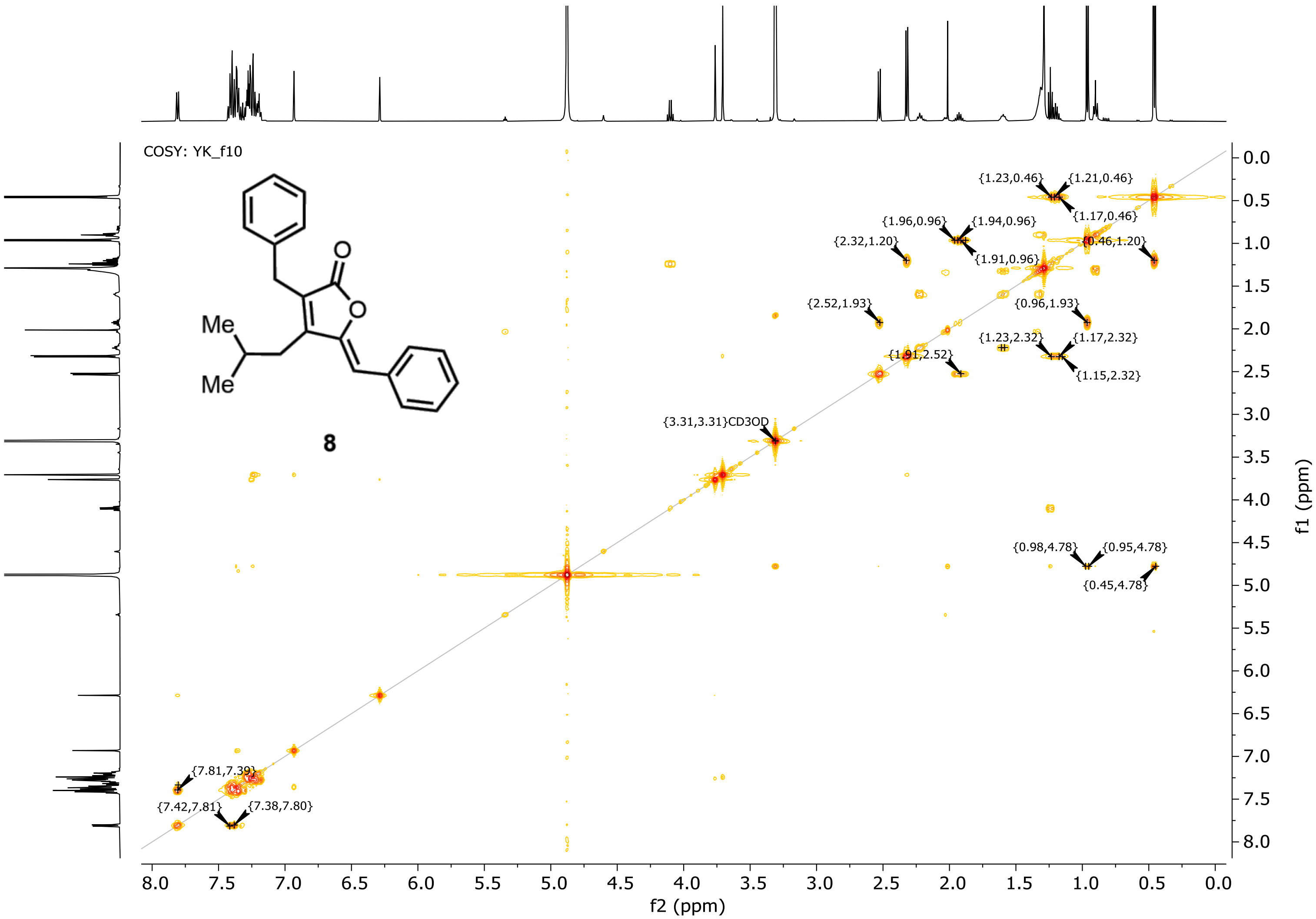


**Figure S38. COSY of deoxyenhygrolide A (8b), measured in MeOD-$d_4$.**

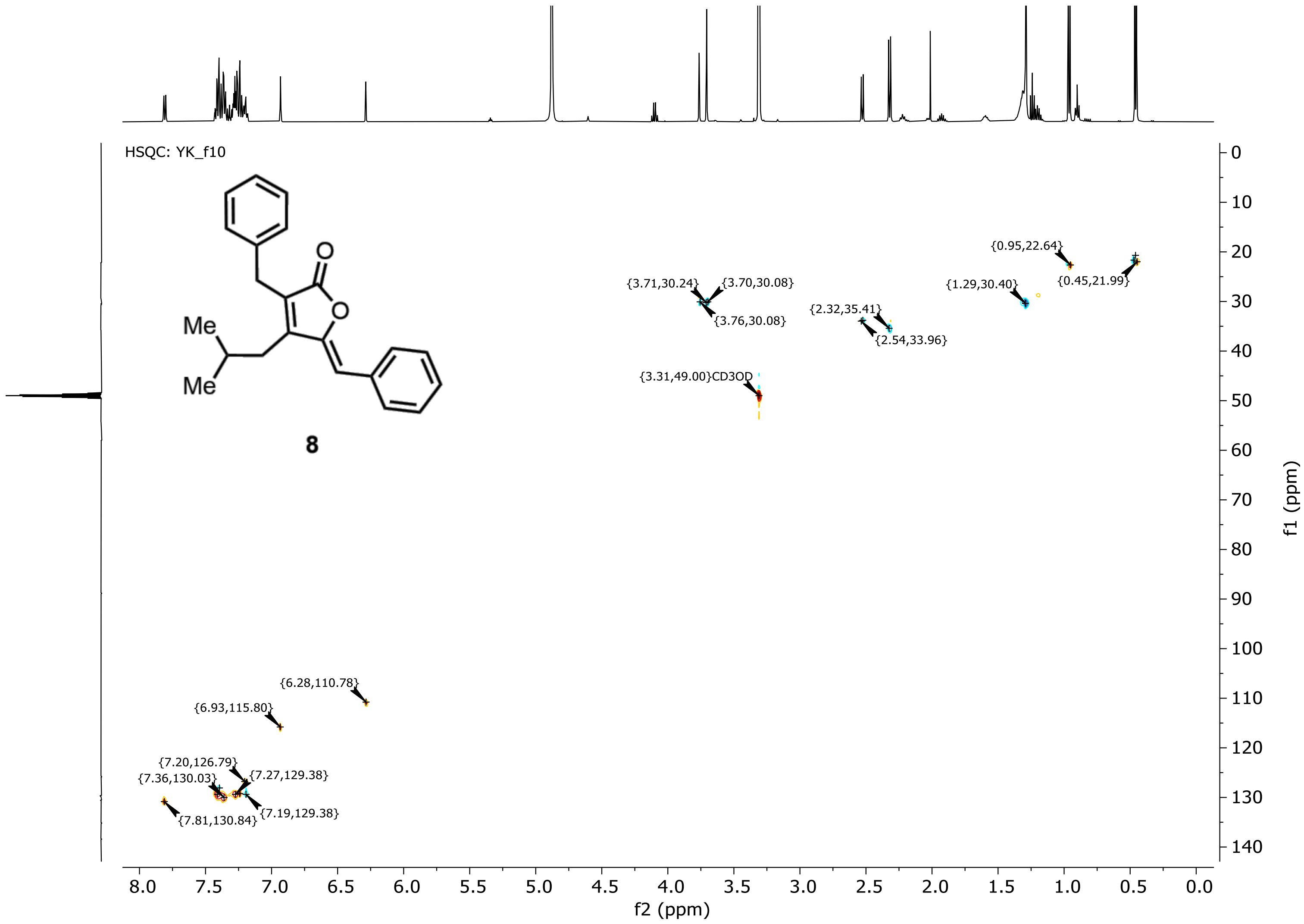


**Figure S39. HSQC of deoxyenhygrolide A (8b), measured in MeOD-$d_4$.**

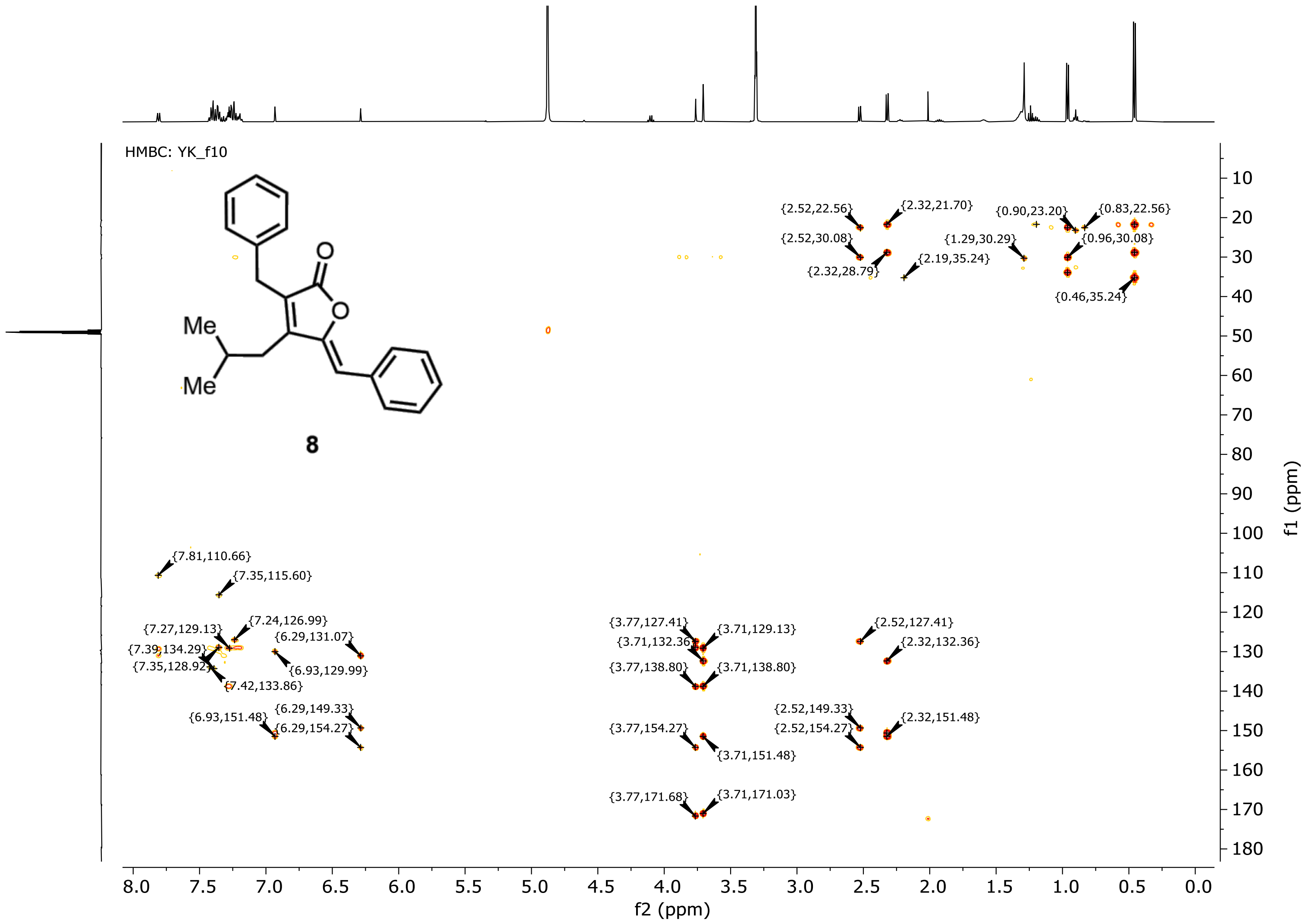


**Figure S40. HMBC of deoxyenhygrolide A (8b), measured in MeOD-d₄.**
